# A pair of readers of histone H3K4 methylation recruit Polycomb repressive complex 2 to regulate photoperiodic flowering

**DOI:** 10.1101/2025.03.23.644783

**Authors:** Xiao Luo, Xueqin Li, Zhijuan Chen, Shu Tian, Yajie Liu, Zhiyun Shang, Jiamu Du, Yuehui He

## Abstract

In flowering plants, the transition from vegetative growth to reproduction or flowering, is often timed by seasonal changes in day length (photoperiod). In the model flowering plant *Arabidopsis thaliana*, the photoperiodic cue increasing day length or long day, through the photoperiod pathway, induces a daily rhythmic activation of the florigen gene *FLOWERING LOCUS T* (*FT*) to promote flowering. Under inductive long days (LDs), *FT* expression is activated around dusk, but to be repressed overnight and into the early afternoon the next day. The mechanism underlying the daily oscillation of *FT* repression to ensure long-day induction of flowering remains unclear. Here, we report that AtING1 and AtING2, Arabidopsis homologs of the mammalian <u>In</u>hibitor of <u>G</u>rowth (ING) proteins, read di- and tri-methylated histone 3 lysine 4 (H3K4me2/me3) on *FT* chromatin and further recruit Polycomb-repressive complex 2 (PRC2) to repress *FT* expression at night and into the early afternoon the next day, following *FT* activation at dusk in LDs. This prevents precocious flowering under inductive LDs. Our study reveals that a previously-undescribed chromatin-regulatory module: H3K4me2/me3-ING1/2-PRC2, which timely represses *FT* expression following the daily rhythmic *FT* activation at dusk by the long-day pathway, to prevent excessive *FT* expression and thus precisely control the timing of the transition to flowering in response to inductive photoperiodic signals.

## Introduction

The timing of developmental transition from vegetative growth to reproduction is crucial for reproductive success in flowering plants. This transition, namely flowering, is regulated by endogenous factors and environmental signals such as seasonal changes in day length (photoperiod)^1,2^. The photoperiod pathway integrates day-length signals with the endogenous circadian clock system to promote the expression of a primary transcription factor that activates the expression of the major florigen genes *FT* (in Arabidopsis) or *FT* homologs (in other flowering plants), resulting in floral induction^1–3^. In the long-day plant Arabidopsis, inductive long day signals induce the production of the zinc-finger transcription factor CONSTANS (CO) that activates *FT* expression to promote flowering^2^.

The expression of the florigen gene *FT* is activated in leaf veins by the CO protein upon the perception of light signals by photoreceptors in leaves, and the FT protein is subsequently transported from the veins to the shoot apical meristem (SAM), to promote the transition from vegetative growth to reproduction at the shoot apex^4–7^. Through the long-day photoperiod pathway, *CO* mRNA expression is gradually upregulated towards the end of the day, and the coincidence of light-dependent CO protein stabilization with the high level of *CO* mRNA in late afternoon results in a gradual accumulation of the CO protein from late afternoon towards dusk (the end of light period)^1,2^. CO assembles into a trimeric complex with the B and C subunits of Nuclear Factor Y (NF-Y), CO-NF, which recognizes *CO*-responsive elements (COREs) located in the proximal promoter of *FT* to activate *FT* expression^8–10^. In addition, a trimeric NF-Y complex, specifically binds to a distal enhancer with a CCAAT motif in the *FT* promoter^8,11^, to facilitate promoter looping between the distal enhancer and proximal COREs, leading to a further activation of *FT* expression under LDs^12–14^. Thus, upon the accumulation of CO towards dusk under inductive LDs, *FT* expression is strongly activated around dusk^1^. The abundance of the CO protein peaks at dusk, but is rapidly degraded by proteasomes at night^1,2^. Interestingly, *FT* expression is swiftly turned off after dusk and remains to be repressed into the early afternoon the next day^15,16^. To date, the underlying mechanism for the rapid *FT* repression following CO-mediated *FT* activation around dusk remains elusive.

Chromatin modifications play an important role to regulate *FT* expression. *FT* chromatin is in a bivalent state bearing both active H3K4me3 and repressive histone 3 lysine-27 trimethylation (H3K27me3)^17^. The H3K27 methyltransferase complex PRC2, composed of the H3K27 methyltransferase CURLY LEAF (CLF) and three core structural subunit including FERTILIZATION INDEPENDENT ENDOSPERM (FIE), EMBRYONIC FLOWER 2 (EMF2) and MULTICOPY SUPPRESSOR OF IRA1^18^, binds to *FT* chromatin to deposit the repressive H3K27me3 in the *FT* promoter as well as gene body, leading to transcriptional repression^14,19^. Upon CO accumulation around dusk in LDs, CO-NF antagonizes CLF-PRC2 on the *FT* promoter chromatin, leading to a reduction of PRC2 enrichment to relieve Polycomb-mediated repression^14^. Furthermore, CO interacts with the chromatin-remodeling factor PICKLE (PKL) that further recruits the H3K4 methyltransferase ARABIDOPSIS TRITHORAX-RELATED PROTEIN 1 (ATX1) to deposit active H3K4me3 and H3K4me2 on *FT* chromatin, and thus promote *FT* activation^20,21^. In addition, CO recruits a pair of chromatin readers MORF-RELATED GENE 1 (MRG1) and MRG2, recognizing both H3K4me3 and histone 3 lysine-36 trimethylation (H3K36me3), to promote *FT* expression around dusk in LDs^22,23^. In addition to histone methylation, histone acetylation and de-acetylation are involved in *FT* regulation^23–25^. Thus, *FT* expression is under sophisticated regulation by various chromatin modifiers.

The ING family proteins, originally named for their function as tumor suppressors in humans, are widely found in fungi, plants and animals^26–28^. ING proteins are composed of an N-terminal coiled-coil domain and a C-terminal plant homeodomain (PHD) zinc finger^27,29,30^. In mammalian cells, ING1 and ING2, through their PHD domains, bind strongly to H3K4me3, but with a lower affinity to H3K4me2, and further recruits the repressive histone deacetylase (HDAC) complex mSin3a-HDAC1 to repressive gene expression^29,31,32^, whereas other mammalian ING proteins including ING3, ING4 and ING5, have been found to associate with histone acetyltransferase (HAT) complexes to activate gene expression^30,33^. In fungi, an ING1 homolog (Fusarium ING-like Protein 1) interacts with a HAT complex to promote gene expression^34^. Two ING proteins including AtING1 and AtING2 have been identified in Arabidopsis, and *in vitro* peptide pulldown experiments have revealed that these proteins can bind an H3K4me3 peptide^27,35^. A further study reveals that *AtING1* is involved in the control of plant height and branch growth^35^. While the mammalian homologs of AtING1/2 have been shown to mediate histone deacetylation and repress gene expression, whether and how plant ING proteins may repress gene expression are unknown.

In this study, we first report that the crystal structures of AtING1 and AtING2 in complex with H3K4me3. Structural and biochemical analyses show that both readers strongly bind both H3K4me2 and H3K4me3. We further found that *AtING1* and *AtING2* function in partial redundancy to repress the floral transition specifically in inductive LDs, but not in short days (SDs). At dusk during a LD cycle, the CO-NF transcriptional activation complex promotes the enrichments of H3K4me2/me3 and further activates *FT* expression. After dusk, CO is degraded, and AtING1 and AtING2 read H3K4me2/me3 on *FT* chromatin and further recruit the CLF-PRC2 H3K27 methyltransferase complex to rapidly repress *FT* expression at night and into the early afternoon the next day, to prevent precocious flowering, in response to inductive LDs. These findings together show that the chromatin-regulatory module of H3K4me2/me3-ING1/ING2-PRC2 prevents excessive *FT* expression and precocious flowering, and functions together with the LD-induced CO accumulation to confer a daily rhythmic pattern of *FT* activation and repression, thereby precisely controlling the timing of the floral transition, in response to inductive LD signals.

## Results

### AtING1 and AtING2 binds strongly to H3K4me2/me3 through their PHD domains

To explore the molecular functions of ING proteins in plants, we first measured the affinity of the PHD finger domains of both AtING1 and AtING2 (PHD^ING1^ and PHD^ING2^) for methylated H3K4 peptides (Figure 1a, b and Figure S1a), using isothermal titration calorimetry (ITC) with purified PHD fingers. The PHD^ING1^ binds to H3K4me2 and H3K4me3 with high affinities of 1.21 μM and 0.86 μM, respectively (Figure 1a). Similarly, the PHD^ING2^ binds to H3K4me2 and H3K4me3 with high affinities of 4.37 μM and 2.91 μM, respectively (Figure 1b). Both PHD^ING1^ and PHD^ING2^ exhibit a high affinity for H3K4me3, comparable to the binding affinity of 1.5 μM observed between the human ING2 PHD finger and the H3K4me3 peptide^29^. In addition, we found that both PHD^ING1^ and PHD^ING2^ can bind to H3K4me1 with weaker affinities of 5.65 μM and 8.70 μM, respectively, but do not bind to unmethylated H3K4 peptides (Figure 1a, b). Thus, the Arabidopsis ING proteins are readers of methylated H3K4 with preference for the higher methylation levels.

**Figure 1.**
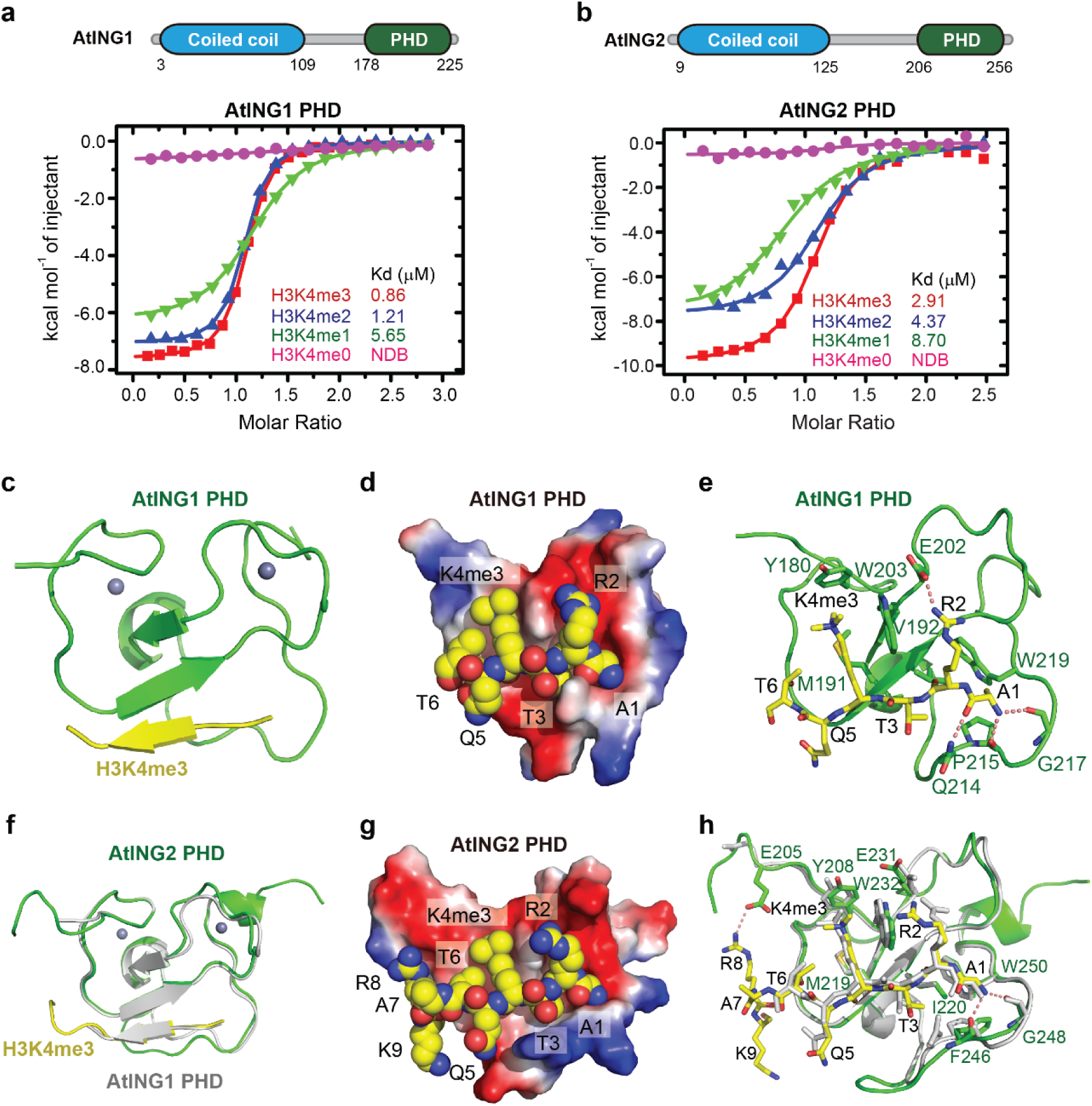
Structural analysis of the binding of PHD fingers of AtING1 and AtING2 to methylated H3K4. **a**, **b**, The isothermal titration calorimetry (ITC) binding curves for AtING1 PHD finger (**a**) and AtING2 PHD finger (**b**) were analyzed in relation to various H3K4 methylation peptides. On top are schematic representations of the domain architecture of AtING1 (**a**) and AtING2 (**b**), with the numbers indicating the amino acid positions of the respective domains. **c**, Overall structure of AtING1-PHD finger in complex with an H3K4me3 peptide in a ribbon diagram with the PHD finger colored in green and the peptide in yellow. The two Zn^2+^ ions are highlighted as gray balls. **d**, An electrostatics surface view of the AtING1-PHD finger with the H3K4me3 peptide in a space filling model. The peptide lies in a negatively charged surface cleft of the PHD finger. **e**, The detailed interaction between the AtING1-PHD finger and the H3K4me3 peptide. The interacting residues are highlighted in stick model and the hydrogen bonds are highlighted as dashed red lines. The H3K4me3 side chain is specifically accommodated by an aromatic cage. **f**, The superposition of AtING2 PHD-H3K4me3 complex (in green and yellow) and AtING1 PHD-H3K4me3 complex (in silver) shows the two complexes share similar folding and peptide binding mode. **g**, An electrostatics surface view of the AtING2-PHD finger with the bound H3K4me3 peptide in space filling model. The peptide attaches to a negatively charged surface cleft of the PHD finger. **h**, The detailed interaction between the AtING2-PHD finger (in green) and the H3K4me3 peptide (in yellow). The interacting residues are shown in stick model and the hydrogen bonds are highlighted as dashed red lines. A superposition of the AtING1-H3K4me3 complex is shown in silver with the interacting residues highlighted in silver stick.

### Crystal structures of AtING1 and AtING2 in complex with H3K4me3

To further investigate the molecular mechanism of recognition of H3K4me2/3 by the Arabidopsis ING PHD fingers, we first determined the crystal structures of PHD^ING1^ in complex an H3K4me3 histone peptide using single-wavelength dispersion (SAD) and refined to a resolution of 1.7 Å, yielding an *R* factor of 0.157 and a free *R* factor of 0.200 (Figure 1c and Table S1). The asymmetric unit contains one PHD^ING1^-H3K4me3 complex. The PHD finger adopts a canonical fold characterized by long loops stabilized by two Zn^2+^ ions, a double-stranded antiparallel β-sheet, and a short α-helix^36^ (Figure 1c). The overall structure of the PHD^ING1^ closely resembles other reported ING family PHD finger structures, particularly the human ING1 PHD finger, with an RMSD of 0.6 Å for 50 aligned Cαs^29,37,38^. The H3K4me3 peptide exhibits good electron density, traceable from Ala1 to Thr6, although Gln5 and Thr6 show relatively poor density in their side chains (Figure S2a). The peptide adopts a β-strand-like extended conformation that pairs with the anti-parallel β-sheet of the PHD finger, forming a continuous three-stranded β-sheet and establishing main chain hydrogen-bonding interactions (Figures 1c and S2b).

Similar to other ING family PHD fingers^29,37,38^, the AtING1 PHD finger features a negatively charged surface cleft that accommodates the H3K4me3 peptide (Figure 1d). The N-terminal amino protons of the H3K4me3 peptide form two hydrogen bonds with the main chain carbonyls of Pro215 and Gly217, effectively docking the N-terminus of the H3 tail (Figure 1e). The methyl group of H3A1 is accommodated by a hydrophobic pocket formed by Val192, Pro215, and Trp219 (Figure 1e). Additionally, the main chain carbonyl of H3A1 forms a hydrogen bond with Gln214 (Figure 1e).

The main chain of N-terminal H3 peptide, from H3R2 to H3T6, pairs with the main chain of Gly189 to Ala193 of the PHD finger, forming an anti-parallel β-sheet (Figure S2b). The side chain of H3R2 engages in hydrogen bonding and salt bridge interactions with Glu202 (Figure 1e). Furthermore, Met191 contributes to the hydrophobic pocket alongside the aromatic cage residues Tyr180 and Trp203, enhancing the hydrophobic recognition of H3K4me3 (Figure 1e). This supports the notion that the AtING1 PHD finger preferentially recognizes the high-level methylation states of H3K4 (Figure 1a).

In addition to AtING1, we further determined the crystal structure of PHD^ING2^ in complex with an H3K4me3 peptide at 1.6 Å solution, yielding an *R* factor of 0.156 and a free *R* factor of 0.199 (Figure 1f and Table S1). AtING1 and AtING2 PHD fingers share a sequence identity of around 60% (Figure S1a), while their structures have an RMSD of 0.93 Å for 50 aligned Cαs upon superposition, indicating almost identical structures (Figure 1f). The H3K4me3 peptide possesses good electron density and can be traced from Ala1 to Lys9, which is longer than observed in PHD^ING1^-H3K4me3 complex (Figure S2c). Generally, the PHD^ING2^ recognizes H3K4me3 peptide using a similar strategy, which employs a negatively charged surface cleft to accommodate the peptide, especially the docking of the N-terminal amino group by a surface pocket and the specific recognition of the tri-methyl lysine by an aromatic cage (Figure 1g, h). PHD^ING1^ and PHD^ING2^ share a similar H3K4me3 recognition mechanism with minor differences in the recognition of H3R2 and H3R8 (Figure 1h). In particular, due to the conformational change of PHD^ING2^ Glu231, the side chain of H3R2 does not form hydrogen bonding and salt bridge interactions with PHD^ING2^ Glu231, while H3R2 forms hydrogen bonds and salt bridge interactions with the equivalent PHD^ING1^ Glu202 in the PHD^ING1^-H3K4me3 complex (Figure 1h). In addition, the H3R8 forms hydrogen bond and salt bridge interactions with Glu205 of PHD^ING2^, while it is disordered in the PHD^ING1^-H3K4me3 complex (Figure 1h).

Collectively, our findings show that AtING1 and AtING2 have high affinities for H3K4me2/me3, with structural analyses revealing similar recognition mechanisms for the H3K4me3 peptide. The crystal structures of ING PHD fingers in complex with H3K4me3 demonstrate a conserved binding strategy involving a negatively charged cleft and an aromatic cage (Figure S2d). This suggests that both proteins can partially compensate for each other’s functions through their shared ability to recognize H3K4me2/me3.

### *AtING1* and *AtING2* act in partial redundancy to specifically repress long-day induction of flowering

To investigate the biological functions of *AtING1* and *AtING2*, we searched for loss-of-function mutants in *ING1* and *ING2* from several Arabidopsis mutant populations^39,40^, and identified an *ing1* mutant with a T-DNA insertion in the third exon^35^, and an *ing2* mutant with a T-DNA insertion in the fifth exon (Figure 2a). These insertions resulted in undetectable full-length transcripts of both genes (Figure S1b, c), suggesting that both mutants are strong loss-of-function alleles. The *ing1* mutant flowered moderately earlier than the wild-type Col-0 (WT), whereas *ing2* flowered only slightly earlier than WT under LD conditions, as measured by the developmental criterion: total number of leaves formed prior to flowering (Figure 2b, c). Next, we constructed an *ing1 ing2* double mutant, and further found that it flowered much earlier than WT or either single mutant in LDs (Figure 2b, c). To confirm that the early flowering phenotype of the *ing1 ing2* double mutant is indeed caused by the loss-of-function mutations in *ing1* and *ing2*, we introduced a wild-type genomic fragment of *ING1* or *ING2* into *ing1 ing2*, and found that the early flowering was fully rescued to that of the single mutants (Figure 2c). Taken together, these results show that *ING1* and *ING2* act in partial redundancy to repress the floral transition in Arabidopsis.

**Figure 2.**
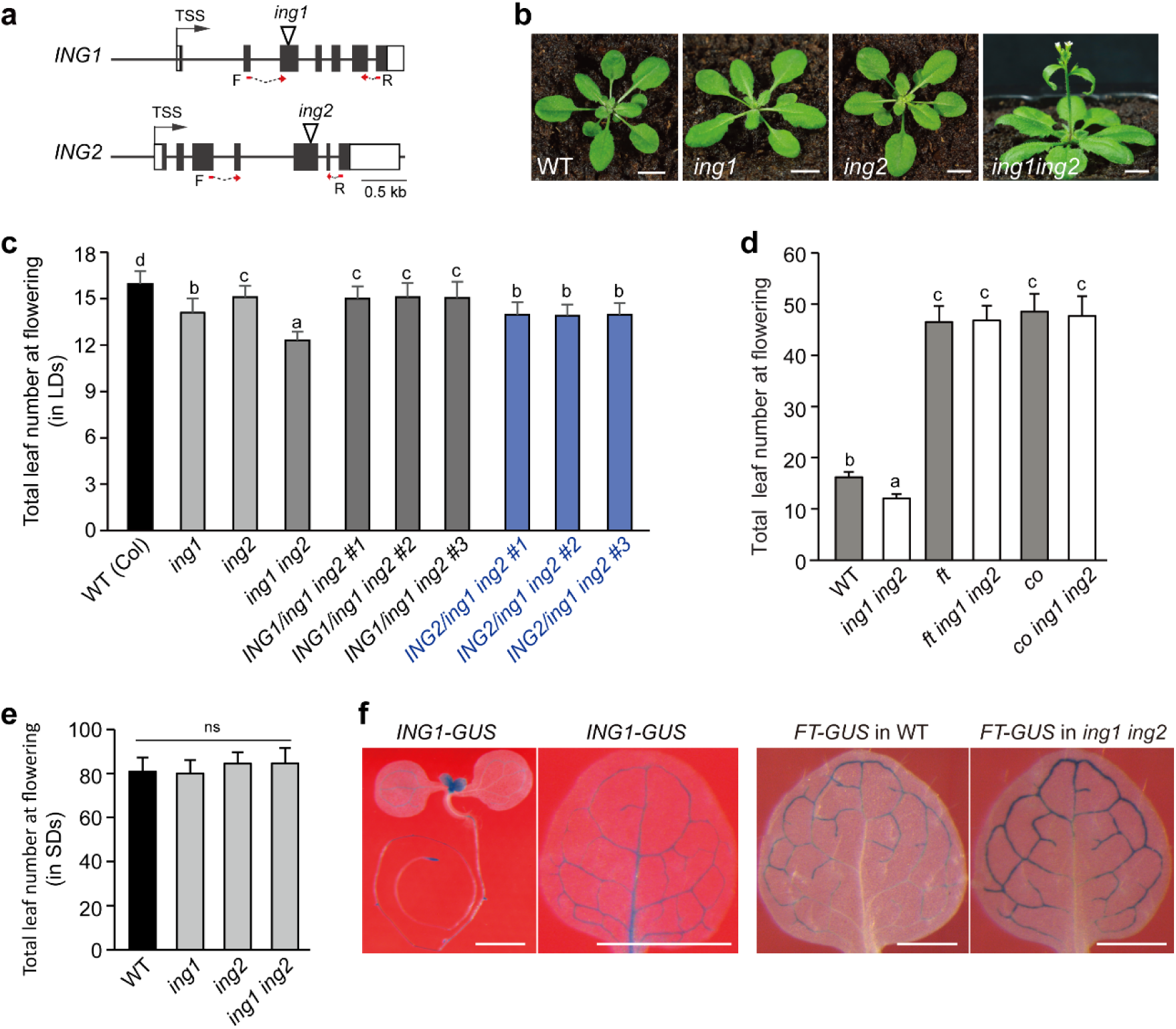
Arabidopsis *ING1* and *ING2* function in partial redundancy to inhibit the floral transition by repressing *FT* expression in LDs. **a**, Schematic representations of *ING1* and *ING2* structures. Filled boxes for coding exons, unfilled boxes for UTR regions, TSS for transcription start site, and inverted triangles for T-DNA insertions. Red lines with arrows show the positions of primers for RT-PCR analysis of *ING1* and *ING2* expression (shown in Figure S1b, c). **b**, Phenotypes of the WT (Col-0), *ing1*, *ing2*, and *ing1 ing2* mutants grown under LDs. Scale bars: 1.0 cm. **c**, Flowering times, represented by the total number of leaves formed before flowering, were scored for *ing1*, *ing2*, *ing1 ing2*, and indicated rescue lines grown in LDs. **d**, **e**, Flowering times of the indicated genotypes grown in LDs (**d**) and SDs (**e**). **f**, Histochemical GUS staining of *ING1-GUS* and *FT-GUS* reporter lines in WT and *ing1 ing2*. Leaves from 10 to 14-d-old seedlings. Scale bars: 1 mm. **c-e**, About 20 plants per line were evaluated. Error bars indicate standard deviation (s.d.). Letters indicate statistically-significant differences (one-way ANOVA, *p*<0.05), ns for not significant.

The developmental transition to flowering in the long-day plant Arabidopsis is known to be induced by increasing day lengths, but inhibited by SDs. Interestingly, unlike in LDs, *ing1*, *ing2* and *ing1 ing2* all flowered like WT under SD conditions (Figure 2e). Thus, *ING1* and *ING2* function specifically to inhibit long-day induction of the floral transition in Arabidopsis. In other words, these *ING* genes function in the long-day photoperiod pathway to regulate flowering.

### *ING1* and *ING2* repress *FT* expression in LDs

The *CO*-*FT* regulatory module is known to mediate long-day flowering regulation^2^; hence, we explored the genetic interactions among *CO*, *FT* and *ING1*/*2*. The *co ing1 ing2* triple mutant flowered as late as *co* (Figure 2d); thus, the *co* mutation is epistatic to *ing1 ing2*. Moreover, the *ft ing1 ing2* mutant flowered as late as *ft*, revealing that *FT* is required for the early flowering upon loss of *ING1* and *ING2* function (Figure 2d). These results show that *ING1* and *ING2* function through the *CO-FT* regulatory module, to regulate the long-day induction of flowering.

Considering that CO activates *FT* expression to induce flowering, we next examined *FT* expression in *ing1*, *ing2*, and *ing1 ing2* seedlings at the end of light period in LDs (ZT16, 16 hour/hr after light on; Zeitgeber time [ZT]). *FT* transcript levels were moderately increased in both *ing1* and *ing2*, but with a more pronounced elevation in the double mutant (Figure 3a), consistent with the partial redundant roles of *ING1* and *ING2* in floral repression. In addition, we found that the level of *FT* expression remained extremely low in both WT and *ing1 ing2* in SDs (Figure 3a). Together, these results reveal that *ING1* and *ING2* function to down-regulate *FT* induction by the LD signals, and thus specifically inhibit the floral transition in LDs, but not in SDs (Figure 2c, e).

**Figure 3.**
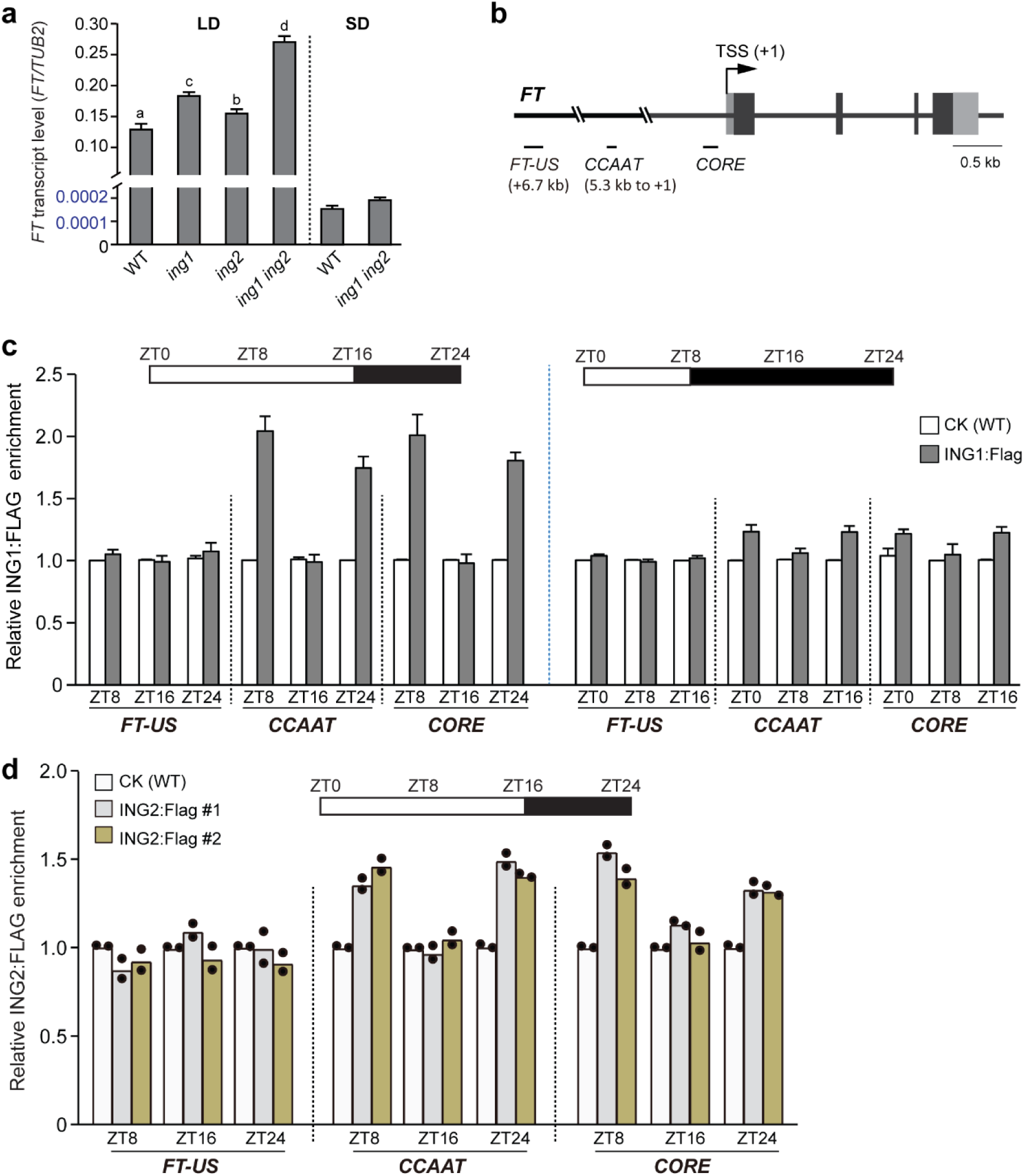
ING1 and ING2 dynamically bind to the distal CCAAT enhancer and proximal CORE regions in *FT* promoter to prevent *FT* overexpression over a LD cycle. **a,** *FT* transcript levels in the indicated seedlings at dusk in LDs (ZT16) and in SDs (ZT8), quantified by RT-qPCR. The levels of *FT* transcript were normalized directly to the endogenous control *TUB2*. Values are means ± s.d. of three biological replicates, and letters indicate statistically significant differences (one-way ANOVA, *p*<0.01). **b**, Schematic representation of the *FT* structure, with filled boxes for exons, and the three regions examined in ChIP-qPCR are indicated. **c**, ChIP-qPCR analysis of dynamic binding of ING1 to *FT* promoter chromatin at the indicated time points over a long-day and short-day cycle. ZT for zeitgeber time (hours after light onset). Levels of anti-FLAG-immunoprecipitated genomic fragments of *FT* were quantified by qPCR, and normalized first to the internal control *TUB2*. Relative ING1:FLAG fold enrichments at three examined *FT* regions in the *ING1:FLAG* line over the background control (WT), are shown. Values are means ± s.d. of three biological replicates. **d**, Dynamic binding of ING2 to *FT* promoter chromatin at the indicated time points over a long-day cycle. Two independent ING2:FLAG lines were analyzed, with two biological replicates conducted for each line. Individual data values are plotted as black dots.

*FT* expression is specifically induced in leaf veins by the LD signals^41^. To further explore the spatial regulation of *FT* expression by *ING1*, we examined the spatial pattern of *ING1* expression by in frame fusing a genomic fragment of *ING1* (a promoter region plus the full-length coding region) to the *β-GLUCURONIDASE* (*GUS*) reporter gene. Histochemical ING1:GUS staining of transgenic seedlings revealed strong *ING1* expression in the leaf veins as well as in root tips and shoot apices (Figure 2f). Next, we introduced *ing1 ing2* into an *FT_pro_-GUS* reporter line^41^, through genetic crossing, and further found that *FT* expression was upregulated specifically in leaf veins in LDs, upon the loss of *ING1* and *ING2* function (Figure 2f). Thus, *ING1* and *ING2* specifically down-regulate *FT* expression in the leaf vasculature to delay the transition to flowering or prevent precocious flowering under inductive LDs.

### ING1 and ING2 dynamically bind *FT* chromatin over a LD cycle

The *FT* promoter features the distal CCAAT enhancer and the proximal CORE region, both of which are crucial for CO-mediated *FT* activation in LDs^9,12^. To determine whether the ING proteins may directly bind to *FT* promoter chromatin, we performed chromatin immunoprecipitation (ChIP) using a transgenic line expressing a fully-functional ING1:Flag (driven by its native promoter; Figure S3a). Over a LD cycle, ING1 was enriched at both the CCAAT enhancer region and proximal CORE region at midday (ZT8), and the binding was lost at dusk (ZT16), but enriched at *FT* promoter again at night (ZT24) (Figure 3b, c). Thus, the ING1 enrichment on *FT* promoter exhibits a robust oscillation over a LD cycle, and is largely antiphasic to the temporal pattern of the CO protein accumulation and binding to *FT* promoter^10,42^. Interestingly, consistent with neither *ING1* nor *ING2* is involved flowering-time regulation in SDs, ING1 was not enriched on *FT* promoter chromatin under SDs (Figure 3c), revealing that the LD signal is required for ING1 enrichment at the *FT* promoter.

We further determined that ING2 enrichments on *FT* promoter chromatin over a LD cycle, using fully-functional *ING2:Flag* lines (Figure S3a). ING2:Flag, like ING1, was enriched at both the CCAAT enhancer region and proximal CORE region at midday, and the binding was lost at dusk, but enriched at *FT* promoter again at night (ZT24) (Figure 3d). Thus, similar to ING1, the ING2 enrichment on *FT* promoter exhibits a robust oscillation over a LD cycle, consistent with that *ING2* functions in partial redundancy with *ING1* to repress *FT* expression. In addition, we found that the levels of ING2 enrichment on *FT* chromatin at both ZT8 and ZT24 were lower than those of ING1 enrichment, in line with that *ING1* plays a greater role in *FT* repression than *ING2* (Figure 3a, c-d).

### Both H3K4me2 and H3K4me3 are enriched in *FT* promoter over a LD cycle, but not in SDs

We have found that both ING1 and ING2 are enriched at the distal *CCAAT* enhancer and proximal *CORE* region in *FT* promoter in LDs. Given that both proteins are reader of H3K4me2/3, we reasoned that the enrichments of H3K4me2/3 on *FT* chromatin may be required for ING1 and ING2 binding. We conducted ChIP assays with anti-H3K4me2 and anti-H3K4me3, to measure the levels of methylated H3K4 at *FT* over a LD cycle. At ZT8, both H3K4me2 and H3K4me3 were enriched highly in the CORE region and moderately in the distal CCAAT enhancer region (Figure 4a). Upon *FT* activation at ZT16, the levels of H3K4me2/me3 were apparently elevated in both the CCAAT region and CORE region, consistent with that these two marks are linked with *FT* activation, and at night the levels of both marks were reduced (Figure 4a). The enrichments of H3K4me2/me3 on *FT* promoter chromatin at both ZT8 and ZT24 are consistent with that ING1 binds *FT* chromatin through reading these two marks. Interestingly, in contrast to the high levels of H3K4me2/me3 at ZT16, neither ING1 nor ING2 binds to *FT* chromatin, indicating that there is a factor to antagonize ING1 and ING2 binding at this time point.

**Figure 4.**
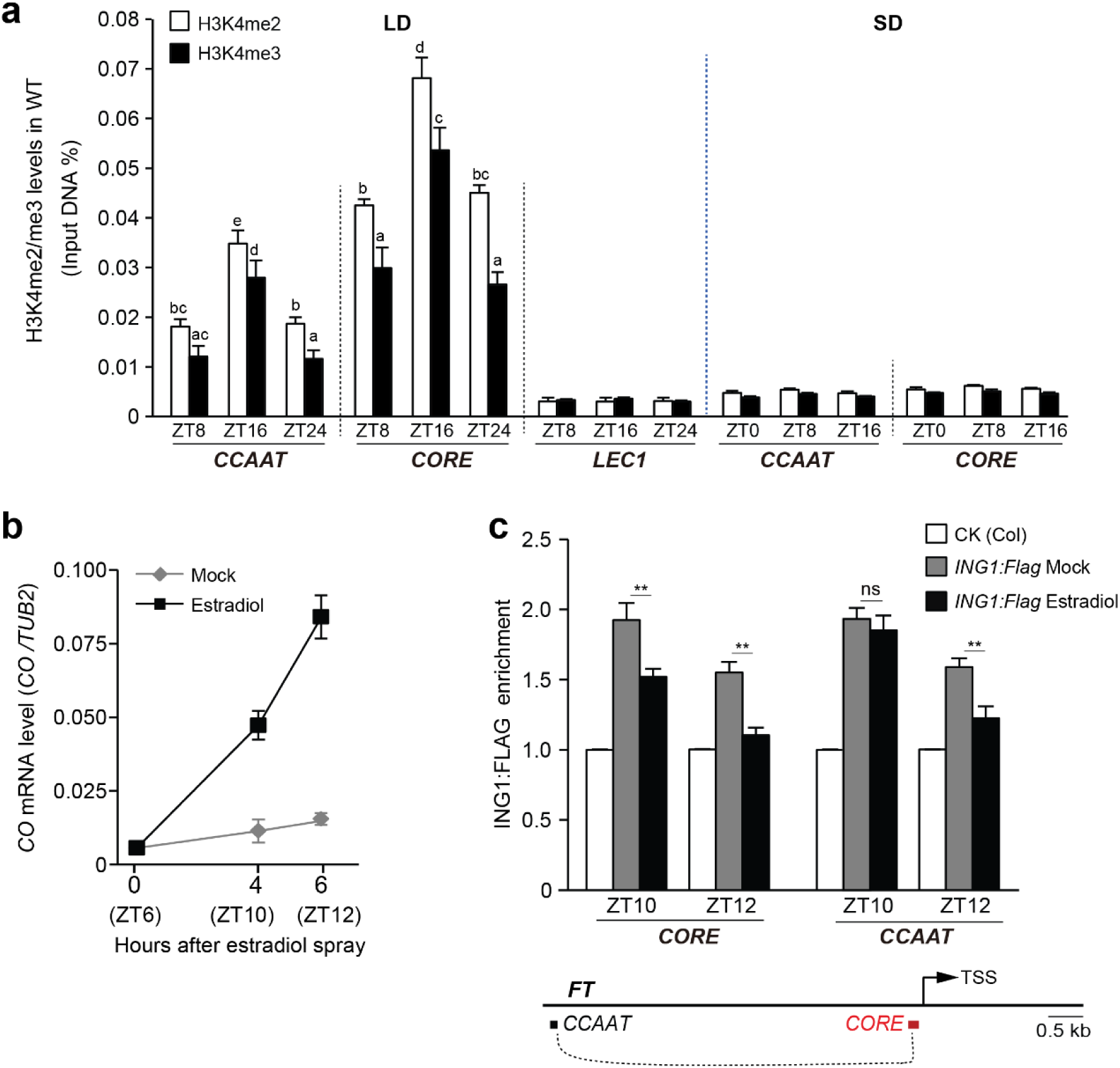
LD-dependent enrichments of H3K4me2/me3 and CO antagonizing ING1 binding *FT* promoter. **a**, ChIP-qPCR measurement of H3K4me2 and H3K4me3 enrichments on *FT* chromatin (in WT) at the indicated time points over a LD cycle and a SD cycle. Levels of the genomic fragments immunoprecipitated using anti-H3K4me3 or anti-H3K4me2, were quantified and normalized directly to ‘input DNA’. The seed-specific *LEC1* (for *LEAFY COTYLEDON 1*, silenced at seedling stages^71^), serves as a negative control lacking of H3K4me2/me3. **b**, *CO* induction line by β-estradiol in an *XVE LexA_pro_-CO* line. Seedlings grown in LDs were treated with either 20 μM estradiol or a mock solution (0.2% ethanol) at ZT6. *CO* levels were assessed at the indicated time points using RT-qPCR and normalized directly to *TUB2*. **c**, ChIP analysis of ING1:FLAG enrichments in *FT* promoter after estradiol application. The doubly hemizygous *LexA_pro_-CO ING1:FLAG* seedlings (F_1_), treated with estradiol or a mock solution at ZT6, were harvested at ZT10 and ZT12. Relative fold enrichments of ING1:FLAG in the examined regions in estradiol- or mock-treated seedlings over a background control (WT /Col), are shown. **a-c**, Values are means ± s.d. of three biological replicates. Letters in (**a**) indicate statistically-significant differences (one-way ANOVA, *p*<0.05), and two-tailed *t* tests were conducted in (**c**), ** *p*< 0.01 and ns for not significant.

We further examined H3K4 methylation on *FT* chromatin in SDs, and found that the levels of both H3K4me2 and H3K4me3 were very low, nearly undetectable (Figure 4a). This is well consistent with that ING1 does not bind to *FT* chromatin under SDs. Together, these results suggest that LD-mediated H3K4me2/me3 enrichments recruit ING1 and ING2 to *FT* promoter chromatin to mediate *FT* repression.

### Recognition of methyl H3K4 is essential for ING1 and ING2 function

To determine ING1 and ING2 bind target chromatin through reading H3K4me2/me3, we first screened the amino acid residues involved in H3K4me2/me3 binding in the PHD fingers of ING1 and ING2, and identified that the Trp203 in ING1 and Trp232 in ING2, located in the aromatic cages for H3K4me2/me3 binding, are essential for methyl H3K4 binding, as the mutations of W203A and W232A eliminated ING1 and ING2 binding H3K4me3, respectively (Figure 5a, b). Next, we examined whether ING1 binding to *FT* chromatin depends on its reading H3K4me2/me3 using the transgenic lines of *ING1:Flag* and *ING1^W203A^:Flag*. In contrast to ING1:Flag, ING1^W203A^:Flag was unable to bind *FT* promoter chromatin at ZT8 in LDs (Figure 5c). These data together reveal that ING1 binds to *FT* chromatin through its reading H3K4me2/me3.

**Figure 5.**
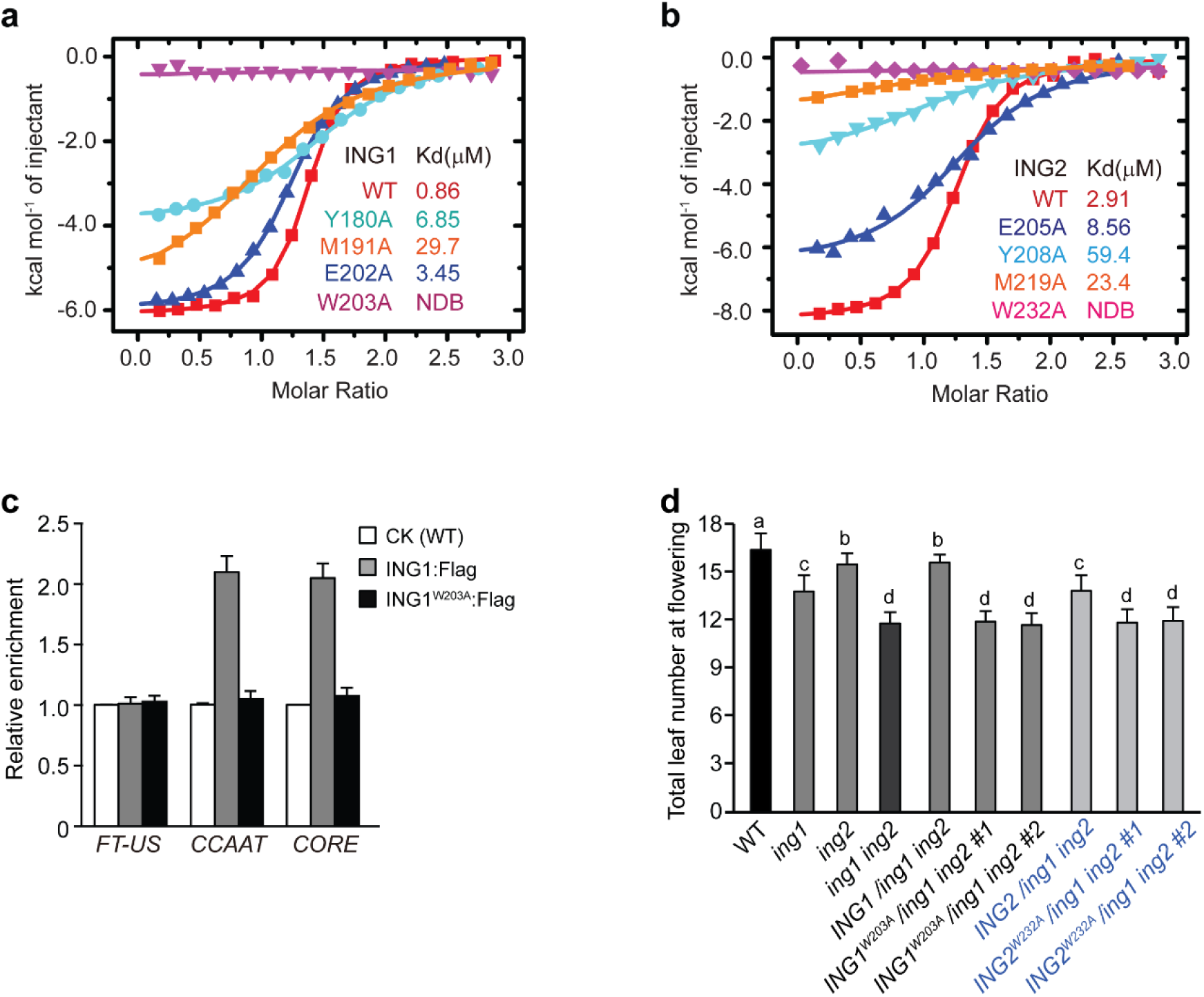
Binding of ING1 and ING2 to methylated H3K4 is essential for their biological functions. **a**, **b**, ITC analysis of the effects of mutated aromatic-cage residues of AtING1 (**a**) and AtING2 PHD fingers (**b**) on H3K4me3 recognition. **c**, ChIP analysis of the bindings of ING1:Flag and ING1^W203A^:Flag to *FT* promoter chromatin. Seedlings expressing *ING1pro-ING1:Flag* or *ING1pro-ING1^W203A^:Flag* (in *ing1 ing2*) were grown in LDs and harvested at ZT8. Anti-Flag-immunoprecipitated DNA fragments from WT serve a background control for fold enrichment calculation. Values are means ± s.d. of three biological replicates. **d**, Flowering times of the indicated lines grown in LDs. *ING1_pro_-ING1^W203A^* and *ING2_pro_-ING2^W232A^* were introduced into *ing1 ing2*. Approximately 20 plants for each line were scored. Bars indicate s.d., and letters denote statistically-significant differences (one-way ANOVA, *p*<0.05).

We further determined whether the biological functions of ING1 and ING2 depend on their reading H3K4me2/me3 on target gene chromatin. First, we introduced *ING1^W203A^:Flag* and *ING2^W232A^:Flag* into the *ing1 ing2* double mutant. Although ING1:Flag and ING1^W203A^:Flag were expressed at similar levels in three examined transgenic lines for each transgene (Figure S3b), only ING1:Flag, but not ING1^W203A^:Flag, was able to rescue the early flowering in *ing1 ing2* (Figure 5d). Similarly, *ING2^W232A^:Flag*, unlike *ING2:Flag*, failed to rescue the early flowering in *ing1 ing2* (Figure 5d). Thus, the recognition of methyl H3K4 by the PHD finger domains from ING1 and ING2 is essential for their functions.

### ING1 binding to *FT* promoters is antagonized by the CO protein

We have found that ING1 binds to *FT* promoter chromatin through its reading H3K4me2/me3; however, despite the high levels of both H3K4me2 and H3K4me3 at ZT16 under LDs, ING1 binding to *FT* chromatin is fully lost (Figure 3c). Given that the CO protein accumulates towards dusk, followed by rapid degradation at night^42^, we reasoned that the CO protein (CO-NF trimeric complex) bound to *FT* promoter antagonizes ING1 (and ING2) binding to *FT* promoter chromatin. Previously, we constructed a chemical induction system of *CO* expression^14^, in which *CO* expression was driven by a *LexA* promoter that is recognized by the chimerical fusion transcription factor XVE (a fusion of the LexA DNA-binding domain, VP16 activation domain and the estrogen receptor^43^). This induction system was introduced into the *ING1:Flag* line by genetic crossing. Subsequently, estradiol was sprayed to the F_1_ seedlings (*LexApro-CO ING1:Flag*), followed by ChIP examination of ING1 binding to *FT* promoter chromatin. Four hr after estradiol application for *CO* mRNA induction, we found that ING1 binding was reduced specifically in the CORE region, but not in the distal CCAAT enhancer region (Figure 4b, c), indicating that CO binding to COREs directly antagonizes ING1 binding to CORE chromatin. Longer *CO* induction (6 hr) caused a reduction of ING1 binding the CCAAT region and further reduced its binding to the CORE region, which led to ectopic *FT* activation (Figures 4c and S4a). The CO-NF complex bound the CORE region facilitates the *FT* promoter looping between the CCAAT distal enhancer and the proximal CORE region, which may result in a reduction of ING1 binding to the CCAAT region upon a high level of CO accumulation. Taken together, these results show that the CO protein antagonizes ING1 binding to *FT* promoter chromatin.

Our results thus far show that the rhythmic *FT* activation induced by the LD photoperiod pathway, leads to the enrichments of both H3K4me2 and H3K4me3 on *FT* promoter chromatin, and that ING1 and ING2 bind to *FT* promoter chromatin to repress *FT* expression, through reading the methylated H3K4. In the morning through early afternoon, INGs bind to *FT* promoter. Upon LD-induced CO accumulation towards dusk, CO antagonizes ING1 (and conceptually ING2) binding to *FT* promoter and activates *FT* expression; subsequently at night, CO is rapidly degraded, and INGs bind to *FT* promoter again to inhibit *FT* expression.

### ING1 and ING2 interact with core subunits of the H3K27 methyltransferase complex PRC2

To explore how the chromatin readers ING1 and ING2 may repress *FT* expression, we searched for transcriptional repressors that may partner with INGs. Previously, we and others have found that Polycomb group (PcG) proteins, particularly CLF-PRC2, play an essential role to repress *FT* expression in both LDs and SDs^14,17,44^; hence, we first examined whether ING1 and ING2 may interact with CLF-PRC2 using yeast two-hybrid assays. We found that ING1 interacted with the H3K27 methyltransferase CLF, and that ING2 interacted with CLF as well as the core structural subunits of PRC2 including FIE and EMF2 in yeast cells (Figure 6a, b). Next, we conducted co-immunoprecipitation (co-IP) to confirm the association of ING1 and ING2 with CLF-PRC2 in Arabidopsis seedlings. Total proteins were extracted from the F_1_ seedlings expressing the functional CLF:HA and ING1:Flag, CLF:HA and ING2:Flag (Figure S4b), or EMF2:VENUS^45^ and ING1:Flag, followed by immunoprecipitation. Indeed, ING1 associated with both CLF and EMF2, and ING2 also interacted with CLF in vivo (Figure 6c-e). Thus, both ING1 and ING2 interact with the chromatin repressor CLF-PRC2, suggesting that INGs engage PRC2 for transcriptional repression in plants.

**Figure 6.**
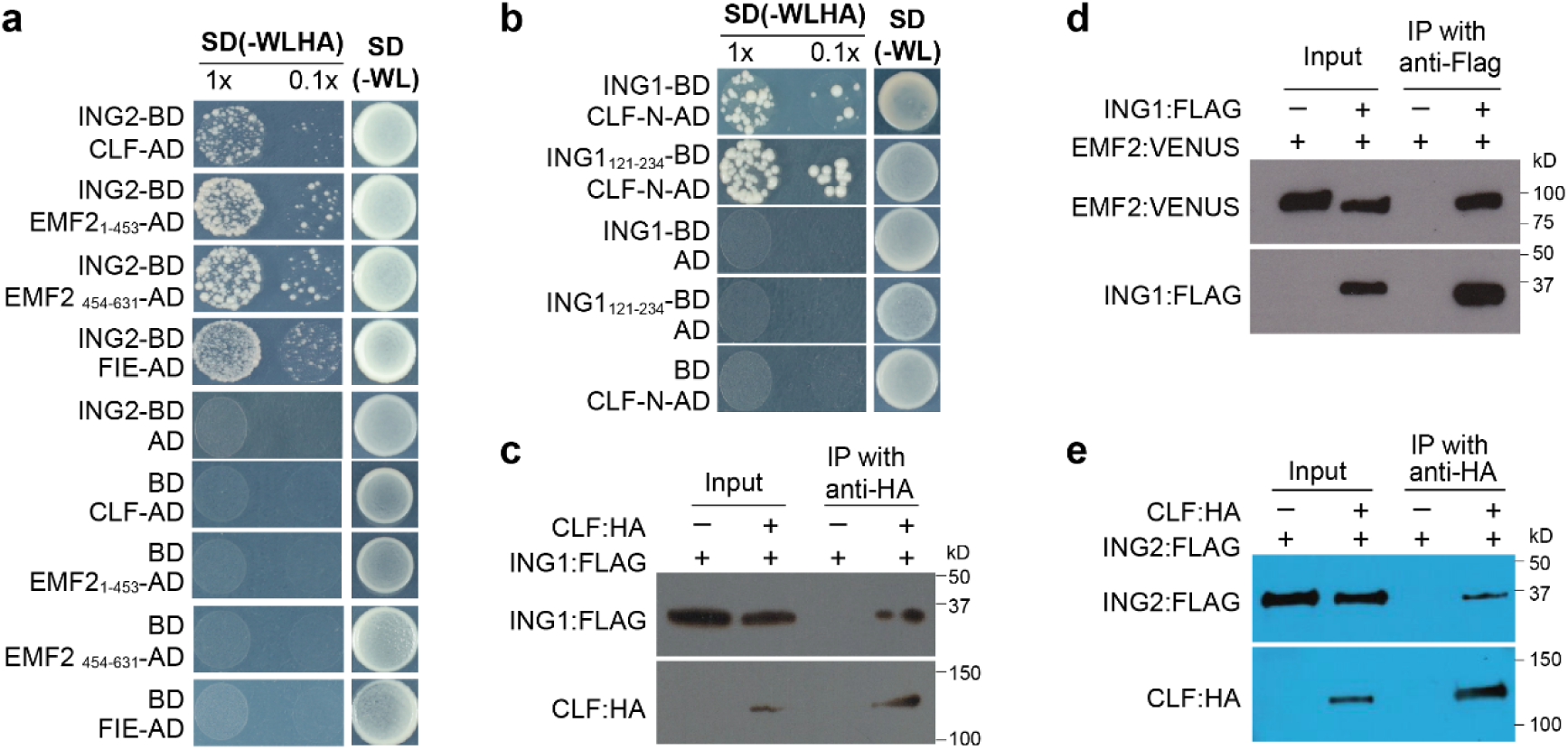
ING proteins interact with CLF-PRC2 in Arabidopsis. **a**, **b**, ING2 interacted with CLF, EMF2 and FIE (**a**), while ING1 interacted with CLF (**b**) in yeast cells. The full-length ING1, ING2, and a N-terminal fragment of ING1 containing the PHD finger (aa 121 to 234) were fused to the GAL4-BD domain. Full-length or fragments of CLF, FIE and EMF2 were fused to GAL4 AD, as indicated. Yeast cells were grown on the stringent selective SD (synthetic defined) media lacking tryptophan (W), leucine (L), histidine (H) and adenine (A), and cells on the non-selective SD media lacking W and L serve as growth controls. **c-e**, Co-immunoprecipitation (co-IP) assays confirming the *in vivo* interaction of ING1:FLAG with CLF:HA (**c**), ING1:FLAG with EMF2:VENUS (**d**), and ING2:FLAG with CLF:HA (**e**). Co-IPs were conducted using total proteins extracted from the F_1_ seedlings expressing paired proteins, followed by western blotting.

### ING1 and ING2 dynamically recruit CLF-PRC2 to mediate repressive H3K27 trimethylation on *FT* chromatin

To determine whether INGs may recruit CLF-PRC2 to *FT* chromatin for transcriptional repression, we conducted ChIP with anti-CLF over a LD cycle in WT and *ing1 ing2* (at a seedling stage). Consistent with previous studies^14^, CLF enrichment on *FT* chromatin exhibited temporal oscillations over a LD cycle: high at ZT8, low at ZT16, and high again at ZT24 (Figure 7d), identical to the temporal pattern of ING1 and ING2 binding to *FT* chromatin (Figure 3c). Upon loss of *ING1* and *ING2* functions, the levels of CLF enrichment at both the CCAAT region and the CORE region were reduced strongly at ZT8, moderately at ZT16, and strongly at ZT24 (Figure 7d). This is consistent with that ING1 and ING2 interact with and dynamically recruit CLF-PRC2 to *FT* chromatin over a LD cycle. Notably, CLF was still partly enriched on *FT* chromatin in *ing1 ing2*, revealing that the recruitment of CLF-PRC2 to *FT* chromatin is partly dependent on ING1 and ING2; in addition, the moderate reduction in CLF enrichment on *FT* promoter chromatin at ZT16 in *ing1 ing2*, compared to WT (Figure 7d), indicates that the CO protein partly antagonizes the ING1- and ING2-independent CLF-PRC2 enrichment at *FT*. There is a possibility that *FT* transcription itself at dusk (ZT16) might cause a reduction in CLF enrichment on *FT* chromatin, but our ChIP examination of CLF enrichment at ZT16 upon the loss of two *FT* repressors known as *AFR1* (for *SAP30 FUNCTION RELATED 1*) and *AFR2*^24^, revealed that the level of CLF enrichment in *FT* promoter remained unchanged (Figure S4c). Taken together, these results show that ING1 and ING2 mediate the dynamic oscillation of CLF-PRC2 on *FT* promoter chromatin over a LD cycle.

**Figure 7.**
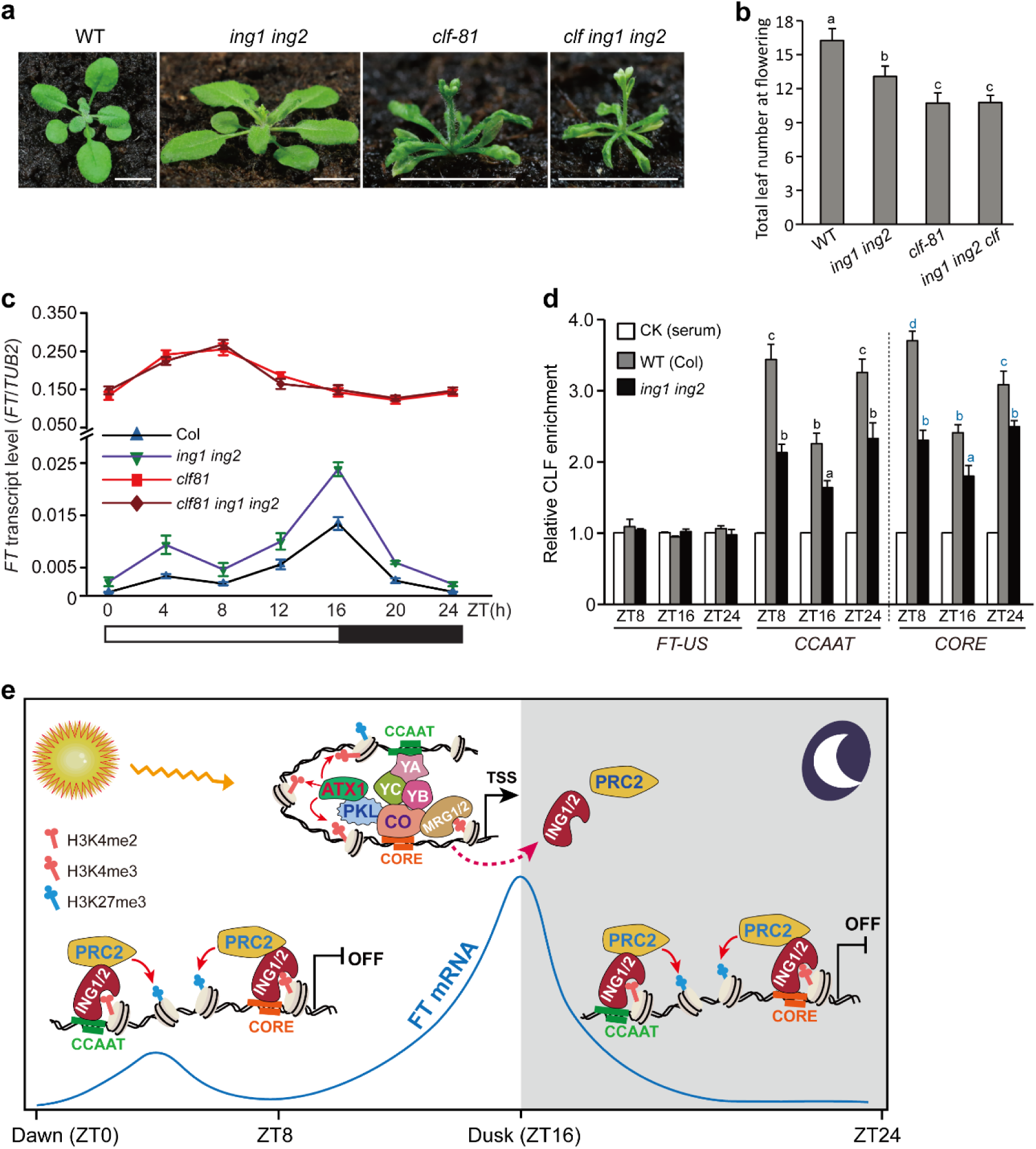
ING proteins mediate dynamic binding of CLF-PRC2 to *FT* promoter chromatin to confer daily rhythmic *FT* repression over a LD cycle. **a,** Phenotypes of *ing1 ing2, clf-81,* and *clf-81 ing1 ing2* grown in LDs. Scale bars: 1.0 cm. **b**, Flowering times of indicated genotypes grown in LDs. About 30 plants were scored for each genotype, and error bars for s.d. **c**, Levels of *FT* transcript in the indicated seedlings over a LD cycle, as revealed by RT-qPCR. **d**, ChIP analysis of the rhythmic binding of CLF to *FT* chromatin in the seedlings of WT and *ing1 ing2* over a LD cycle. Seedlings were harvested at ZT8, ZT16, or ZT24. A rabbit polyclonal anti-CLF^62^ or serum were used for ChIP. Relative fold enrichments of CLF in WT and *ing1 ing2* over the background control (WT IP with rabbit serum), are presented. **c**, **d**, Values are means ± s.d. of three biological replicates. Statistically-significant differences within a group of means in (**d**), are indicated by the letters of the same color (one-way ANOVA, *p*<0.05). **e**, A working model for the photoperiodic regulation of *FT* expression by the CO-H3K4me2/me3-ING-PRC2-*FT* regulatory module. Inductive LDs lead to the CO protein accumulation towards dusk. CO, together with NF-YB and NF-YC, form the CO-NF transcriptional activation complex that binds COREs to antagonize ING binding to CORE chromatin, and through the CO-induced promoter looping between the proximal CORE region and distal CCAAT enhancer, ING binding to the distal enhancer region is reduced. Furthermore, CO recruits the PKL-ATX1 module to promote H3K4 methylation on *FT* promoter chromatin, and further engages the chromatin readers MRG1/2 that recognize H3K4me2/me3 on *FT* promoter chromatin to directly antagonize ING binding to *FT* chromatin. Following *FT* activation at dusk, CO is rapidly degraded, and ING1 and ING2 read the existing H3K4me2/3 on *FT* promoter chromatin and further recruit PRC2 to repress *FT* expression overnight and into the early afternoon the next day, until CO begins to accumulate again towards dusk. Notably, *FT* promoter chromatin is in a bivalent state bearing both methylated H3K4 and H3K27me3 throughout a LD cycle.

CLF-PRC2 deposits H3K27me3 on *FT* chromatin to repress *FT* expression^14,17^. We further measured the changes of H3K27me3 on *FT* promoter chromatin in the seedings of WT and *ing1 ing2* over a LD cycle. Prior to dusk (at ZT8), H3K27me3 in both the CCAAT enhancer region and CORE region were at high levels, and loss of *ING1* and *ING2* function led to a reduction of H3K27me3 (Figure S4d). Upon CO accumulation at dusk, the levels of H3K27me3 on *FT* promoter chromatin were apparently reduced, but at night H3K27me3 increased again, which required the functional *ING1* and *ING2* (Figure S4d). These data are consistent with that ING1 and ING2 recruit CLF-PRC2 prior to and after dusk under inductive LDs, to deposit the repressive H3K27me3 for *FT* repression.

To genetically confirm that the ING proteins repress *FT* expression through CLF-PRC2 under inductive LDs, we first examined genetic interactions between *ing1 ing2* and *clf* in flowering-time regulation, and found that *clf* and the *clf ing1 ing2* triple mutant flowered early and at the same time (Figure 7a, b). Thus, INGs indeed engage CLF-PRC2 to repress the floral transition. Next, we further measured the levels of *FT* transcript in the seedlings of WT, *clf*, *ing1 ing2*, and *clf ing1 ing2* over a LD cycle. Loss of *ING1* and *ING2* function gave rise to *FT* de-repression prior to and after dusk, namely, a largely constitutive *FT* expression in LDs (Figure 7c). Loss of *CLF* function caused a great *FT* depression throughout a LD cycle, and *FT* was expressed at the same levels in *clf* and *clf ing1 ing2* (Figure 7c). Taken together, these results show that ING1 and ING2 recruit CLF-PRC2 prior to and after dusk, to repress *FT* expression and thus prevent precocious flowering under inductive LDs.

## Discussion

Under inductive LDs, the photoperiod pathway output CO rhythmically accumulates towards dusk, which in turn activates the florigen gene *FT* expression daily around dusk, conferring the long-day induction of flowering^1,2^. *FT* expression is swiftly turned off after dusk and remains to be repressed from the coming morning until late afternoon^1,2^. Previous studies suggest that the degradation of the CO protein, occurring at night and into the morning through early afternoon in LDs, leads to a rhythmic *FT* repression^42,46,47^. In this study, we have found that following the CO-mediated *FT* activation around dusk, the H3K4me2/me3-ING1/ING2-PRC2 regulatory module plays a crucial role to repress *FT* expression at night and into the late afternoon of the following day (when CO accumulates again). We show that LD signals, through the photoperiod pathway, promote the enrichments of H3K4me2/me3 on *FT* promoter chromatin. Prior to dusk during a LD cycle, the chromatin readers ING1 and ING2 specifically read H3K4me2/me3 at *FT* and further recruit CLF-PRC2 to repress *FT* expression. Upon its accumulation towards dusk, the CO protein (the CO-NF trimeric transcriptional activation complex) antagonizes the binding of INGs to *FT* promoter chromatin, and further promotes both H3K4 di- and tri-methylation and activates *FT* expression around dusk. After dusk, CO is degraded, and INGs come back to read the H3K4me2/me3 on *FT* promoter chromatin and further recruit CLF-PRC2 to repress *FT* expression. Thus, the chromatin-regulatory module of H3K4me2/me3-ING1/ING2-PRC2 prevents excessive *FT* expression and precocious flowering, and functions together with the LD-induced CO accumulation to confer the daily rhythmic pattern of *FT* activation and repression, thereby precisely controlling the timing of the floral transition, in response to inductive LD signals (Figure 7e). These findings collectively define a novel chromatin-regulatory module that plays an important role in the photoperiodic regulation of flowering in LDs.

In this study, we provide the first structural evidence that the plant homologs of mammalian ING1 and ING2 function as the readers of both H3K4me2 and H3K4me3. We further show that both H3K4me2 and H3K4me3 are enriched on *FT* promoter chromatin in a LD-dependent manner. Previous studies have revealed that upon the CO accumulation around dusk in LDs, CO interacts with and recruits the ATP-dependent chromatin remodeler PKL to *FT* chromatin, and that PKL further recruits the H3K4 methyltransferase ATX1 to deposit the active H3K4me3 (and likely H3K4me2), to promoter *FT* expression^20–22^. In plants, H3K4me3 typically promotes or is associated with active gene expression, and H3K4me2 is often found in genomic regions that are actively transcribed and thus is associated with gene expression^48^, but the levels of H3K4me2 in gene body is negatively correlated with transcription levels^49,50^, partly due to the co-transcriptional demethylation of H3K4me2 by a demethylase associated with a transcription-elongation factor^50^. Interestingly, we have found that both H3K4me2 and H3K4me3 were enriched highly in the proximal promoter region (CORE region) and moderately in distal regions, upon the CO accumulation at dusk in LDs. Furthermore, the levels of H3K4me2/me3 were only reduced, but both marks remain well-preserved after dusk (e.g. ZT24) and into the early afternoon the next day (e.g. ZT8); however, both H4K4me2 and H3K4me3 are nearly undetectable on *FT* promoter chromatin in SDs. Thus, the LD-dependent enrichments of H3K4me2/me3 underlie the dynamic binding of ING1 (and likely ING2) to *FT* promoter chromatin in a LD-dependent manner.

We have found that upon its accumulation around dusk, the CO protein antagonizes the binding of INGs to *FT* promoter chromatin, resulting in a loss of the binding at dusk, to promote *FT* expression. The trimeric CO-NF transcriptional activation complex specifically binds the four C/TCACA motifs the CORE region^10,51^, which conceptually antagonizes the binding of INGs to the CORE region. In addition, CO not only promotes H3K4 trimethylation, through the CO-PKL-ATX1 module, but also interacts with and engages the chromatin readers MRG1 and MRG2 that read H3K4me2/me3 on *FT* promoter chromatin and further engage histone acetyltransferases to promote *FT* activation^22,23^. The binding of MRGs to H3K4me2/me3 is expected to directly antagonize the binding of INGs to H3K4me2/me3 on *FT* promoter chromatin. Taken together, these mechanisms collectively result in the loss of ING1 and ING2 binding to *FT* chromatin.

*FT* expression is under intensive regulation by multiple chromatin modifications. During CO-mediated *FT* activation, H3K4 methylation, H3K36 trimethylation and histone acetylation function in concert to promote *FT* expression^22,23^, whereas histone deacetylation, and H3K27me3 and Polycomb-mediated repression function to repress *FT* expression^14,24,52^. *FT* chromatin has been shown to be in a bivalent state with the co-existence of active H3K4me3 and repressive H3K27me3 on the same nucleosome^17^. The genes in the bivalent state are often poised for transcriptional changes^53^. Consistently, *FT* expression state exhibits daily oscillation under inductive LDs: repressed prior to dusk, activated by CO-NF at dusk, and repressed again after dusk^1^, and underlying this daily oscillation is the rapid changes of *FT* chromatin state over a LD cycle (Figure 4a and Figure S4d)^14,24^. Interestingly, we have found that the enrichments of H3K4me2/me3 on *FT* promoter chromatin around dusk, are not eliminated at night, but are recognized and preserved by INGs to recruit the repressive CLF-PRC2 for *FT* repression until the coming late afternoon. Upon the CO protein accumulation towards dusk, the loss of ING binding releases the existing H3K4me2/me3 on *FT* promoter chromatin to facilitate or boost *FT* activation. Thus, during the long day pathway-mediated daily oscillation of *FT* activation, the existing H3K4me2 and H3K4me3 are partly utilized by INGs to rapidly shut down *FT* expression at night, and are subsequently exploited to promote *FT* activation by CO-NF at the coming dusk (Figure 7e). This mechanism highlights an elegant and complexed regulatory system to modulate photoperiodic control of flowering, in response to day-length signals.

Both H3K4me2 and H3K4me3 are commonly found in transcribed regions in the Arabidopsis genome. In this study, we have revealed a novel chromatin-regulatory module: H3K4me2/me3-ING1/2-PRC2, that represses the expression of target genes with the active mark H3K4me3 in plants. A recent Arabidopsis transcriptomic analysis has revealed that 1,482 genes are differentially regulated upon loss of *ING1* function, among which 1,053 genes are upregulated^35^, consistent with our finding that *ING1* typically functions as a transcriptional repressor. Previous studies have shown that the mammalian homologs of AtING1/2 interact with mSin3a/HDAC1/2 complexes to repress target gene expression^29,37^. It is likely that the plant ING1/2 may recruit HDACs to repress target gene expression, in addition to PRC2. The ING family proteins are also involved in gene activation. Previous studies in both yeast and mammals have shown that INGs can interact with histone acetyltransferase complexes, such as the NuA4 core complex, to activate gene expression^30,33^. Not surprisingly, it has recently been found that AtING1 interacts with a core subunit of a HAT complex to regulate a subset of genes^35^. Thus, the plant INGs can interact with both repressive and active chromatin modifiers to regulate target gene expression.

CO, INGs, PRC2 and various chromatin modifiers involved in *FT* repression are evolutionarily conserved across most of the flowering plants^1,18,27,54^. Hence, the H3K4me2/me3-ING-*FT* regulatory module may be functional in photoperiodic control of flowering in other flowering plants. Interestingly, a recent study has revealed that in the long-day plant *Medicago truncatula*, the *ING2* homolog *MtING2* functions to promote the expression of the *FT*-like gene *MtFTa1* to accelerate the transition to flowering^28^, unlike the Arabidopsis INGs to repress *FT* expression. It is likely that MtING2 might interact with an active chromatin factor such as HAT, to regulate *MtFTa1* expression. Hence, upon their reading of H3K4me2/me3, the plant ING proteins may recruit active or repressive chromatin modifiers to activate or repress the expression of *FT* /*FT*-like genes, to regulate flowering time in response to diverse environmental cues.

## Methods

### Plant materials and growth conditions

All of the mutants and transgenic lines used in this study are in the *A. thaliana* Columbia (Col) background. *ft-10*, *co-9*, *clf-81*, *clf-29*, *afr1 afr2*, the *LexA_pro_-CO* line, the *FT_pro_-GUS* reporter line, and the *EMF2:VENUS* line have been described previously^14,24,41,45,55–58^. The *ing1* (SALK_009598) and *ing2* (GK-166D07) mutants were obtained from the Arabidopsis Biological Resource Center. Seeds were germinated and plants were grown on half-strength MS media (agar plates) or in soil at around 22℃ in LDs (16-hr light/8-hr dark) or SDs (8-hr light/16-hr dark) under cool white fluorescent light.

### Plasmid construction

To construct *ING1_pro_-ING1:GUS*, a 3.1-kb *ING1* genomic fragment (1.0-kb 5’ promoter plus the 2.1-kb full-length genomic coding sequence except the stop codon) was fused in frame with *GUS* in the binary vector *pMDC162* ^59^ using Gateway technology. To create *ING1_pro_-ING1*, a 3.5-kb genomic fragment of *ING1* (1.0-kb 5’ promoter, 2.1-kb genomic coding sequence and 0.4-kb 3’ region), was cloned into the *pHGW* vector^60^ via Gateway technology. To construct *ING2_pro_-ING2*, a 3.5-kb genomic fragments of *ING2* (1.1-kb 5’ promoter, 1.9-kb genomic coding sequence and 0.5-kb 3’ region), was cloned into the *pHGW* vector^60^. Additionally, point mutations were introduced into these fragments using overlap PCR to generate *ING1_pro_-ING1^W203A^* (the tryptophan at position 203 is mutated to alanine), and *ING2_pro_-ING2^W232A^* (the tryptophan at position 232 is mutated to alanine); subsequently, these fragments were cloned into the *pHGW* vector.

To generate *ING1_pro_-ING1:FLAG*, *ING1_pro_-ING1^W203A^:FLAG*, and *ING2_pro_-ING2:FLAG* plasmids, a 3.1-kb genomic fragment of *ING1* or *ING1^W203A^* (1.0-kb 5’ promoter and 2.1-kb genomic coding sequence except the stop codon), or a 3.0-kb *ING2* fragment (1.1-kb promoter and 1.9-kb genomic coding region without the stop codon), was fused individually with a *FLAG* tag (three copies /3x); subsequently, these fusions were cloned into the *pHGW* vector. To generate a HA-tagged CLF fusion, a 6.6-kb genomic *CLF* fragment (comprising 2.1-kb promoter and 4.5-kb coding sequence, excluding the stop codon) was fused with a 9xHA tag and subsequently cloned into the *pBGW* vector.

### Gene expression analysis

Total RNAs were extracted from around 10-d-old seedlings grown in LDs or SDs, using the *RNeasy Plus Mini Kit* (Qiagen), following the manufacturer’s instructions. RNAs were then used as templates for cDNA synthesis with oligo(dT) primers using the M-MLV reverse transcriptase (Promega). Real-time quantitative PCR (qPCR) was performed using a *QuantStudio6 Flex* real-time PCR system (Applied Biosystems). The constitutively-expressed *TUBULIN2* (*TUB2*) gene serves as an internal control for normalization, and the ratio of the transcript level of the gene of interest to *TUB2* was calculated using the 2^-ΔCT^method. Primers are listed in Table S2.

### Yeast two-hybrid assay

Yeast two-hybrid assays were performed according to the instructions from the *Matchmaker GAL4 Two-Hybrid System 3* (Clontech). Plasmid pairs were introduced into the yeast strain *AH109*, and yeast cells were then spotted onto a stringent-selection media lacking leucine (L), tryptophan (W), histidine (H), and adenine (A) to detect protein-protein interactions, while the non-selective SD media lacking leucine and tryptophan served as a growth control.

### Chromatin immunoprecipitation

ChIP experiments were performed using the *Magna ChIP* kit (Millipore), following the manufacturer’s instructions with minor modifications^61^. Briefly, the seedlings of interest were harvested at various time points, followed by total chromatin extraction. Immunoprecipitation was carried out with anti-FLAG (Sigma, M8823), rabbit polyclonal anti-CLF^62^, anti-H3K27me3 (Millipore, 07-449), anti-H3K4me3 (Millipore, 05-745R), or anti-H3K4me2 (Millipore, 07-030). The immunoprecipitated genomic fragments of interest were quantified using qPCR on an *ABI QuantStudio6 Flex* real-time PCR system with a SYBR Green PCR master mix. Three biological replicates were analyzed in each experiment. Primers are listed in Table S2. Relative enrichment of the protein of interest at an examined region was initially normalized to the endogenous control *TUB2*, followed by normalization to a background control (WT).

### Histological β-Glucuronidase activity staining

GUS activity staining was performed as previously described^63^. Briefly, transgenic seedlings grown in LDs were harvested at ZT12 and incubated in 90% acetone at -20°C for around 30 minutes. The seedlings were then stained in a 0.5mg/ml X-Gluc (5-bromo-4-chloro-3-indolyl-β-D-glucuronic acid) solution at 37°C for 3 hr or 12 hr depending on the GUS activity, followed by chlorophyll removal with 70% ethanol. Stained seedlings were examined using a Zeiss dissecting microscope.

### Co-immunoprecipitation

Co-immunoprecipitation (Co-IP) assays were conducted as previously described with minor modifications^64^. Briefly, total proteins were extracted from the F_1_ seedlings expressing *CLF_pro_-CLF:9xHA* and *ING2_pro_-ING2:3x FLAG* (double hemizygote), *CLF_pro_-CLF:9xHA* and *ING1_pro_-ING1:3x FLAG* (double hemizygote) or *EMF2:VENUS* and *ING1_pro_-ING1:3x FLAG* (double hemizygote). Then, immunoprecipitations were conducted with anti-HA affinity gel (Sigma) or anti-FLAG affinity gel (Sigma). The proteins of interest in the precipitates were detected by western blotting using anti-FLAG (Sigma, A8592), anti-HA (Roche, 12013819001) or anti-GFP (abcam, ab290).

### β-estradiol application

β-estradiol application was conducted as previously described^14^. In brief, about 10-day-old seedlings grown on the half-strength MS media were sprayed with 20 µM β-estradiol or a mock solution (0.2% ethanol). β-estradiol (Sigma) was dissolved in ethanol and prepared as a 10 mM stock solution.

### Protein expression and purification

The DNA fragment of the PHD finger of Arabidopsis ING1 (residues 163-234), was cloned into a self-modified *pMAL* vector (New England Biolabs), resulting in a fusion with an N-terminal hexahistidine and the Maltose-Binding Protein (MBP) tag; subsequently, the plasmid was introduced into the *E. coli* strain BL21 (DE3) by transformation. Protein expression was induced by adding 0.2 mM IPTG (isopropyl-β-d-thiogalactoside) to the cell culture once its OD600 reached around 0.7. The recombinant His-MBP-PHD^ING1^ was purified using a nickel column (GE Healthcare), and then the His-MBP tag was digested by a TEV protease and removed by a nickel column. The PHD^ING1^ protein was further purified using a Superdex G75 gel filtration column (GE Healthcare), and concentrated to 30 mg/ml. The CDS of the PHD finger fragment of Arabidopsis ING2 (residues 189-262) was cloned, expressed, and purified using the same procedures as above. All point mutants generated using a PCR-based method, were expressed and purified using the above procedures. All the peptides were purchased from GL Biochem Ltd (Shanghai).

### Crystallization, data collection and structure determination

The purified PHD^ING1^ protein was mixed with the histone peptides of H3K4me3 with a molar ratio of 1:4, and incubated at 4 °C for 30 min. The crystallization was carried out using the hanging drop vapor diffusion method at 20°C, in a condition of 40% PEG400, 5% PEG3000, and 0.1 M MES, pH 6.0. The PHD^ING2^-H3K4me3 complex was crystallized in a condition of 0.1 M NiCl_2_, 20% PEG2000 MME, and 0.1 M Tris-HCl, pH 8.5. All crystals were soaked into the reservoir solution supplemented with 15% glycerol and flash-cooled into liquid nitrogen. The diffraction data for PHD_AtING1_-H3K4me3 complex and PHD_AtING2_-H3K4me3 complex were collected at the beamline BL19U1 and BL18U1 of the National Center for Protein Sciences Shanghai (NCPSS) at the Shanghai Synchrotron Radiation Facility (SSRF)^65^. Subsequently, the data were processed with the program HKL3000^66^, and a summary of the statistics of the data collection is presented in Table S1.

The structure of PHD^ING1^-H3K4me3 complex was solved using the SAD method with the program Phenix^67^. Because of the high resolution, almost all the traceable residues were located by the auto-build function of Phenix. Several rounds of manual model building using the program Coot and structure refinement using the program Phenix, were then conducted^67,68^. The structure of PHD_AtING2_-H3K4me3 complex was determined similarly as the PHD_AtING1_-H3K4me3 complex structure. The structural refinement statistics are summarized in Table S1. The molecular graphics were prepared using the Pymol program (DeLano Scientific LLC), and the sequence alignment of INGs was carried out using the program T-coffee and illustrated using the server ESPrit^69,70^.

### Isothermal titration calorimetry

The binding assays of PHD^ING1^ and PHD^ING2^ to histone H3 peptides were performed on a Microcal PEAQ-ITC instrument (Malvern). Proteins were dialyzed against a buffer of 100 mM NaCl, 20 mM HEPES, pH 7.0, and 2 mM β-mercaptoethanol, and the H3 peptides were dissolved into the same buffer. The titration data were fit using the Origin 7.0 program (OriginLab Corporation).

## Data availability

The coordinates and structure factors have been deposited in the Protein Data Bank under accession codes 9M4R and 9M4S.

## Acknowledgements

We are grateful to the staff members from the beamline BL19U1 and BL18U1 of the National Center for Protein Sciences at Shanghai (NCPSS) at the Shanghai Synchrotron Radiation Facility (SSRF) for their assistance during data collection. We thank Dr. Xiaolin Zeng for assistance in plasmid construction. This work is supported in part by the National Key R&D Program of China (2024YFA1306703 to Y.H.), the National Natural Science Foundation of China (31970327 to X.L. and 32325008 to J.D.), Natural Science Foundation of Shandong Province (ZR2021YQ16 to X.L.), Taishan Scholar Foundation of Shandong Province (tsqn202211301 to X.L.), Shenzhen Science and Technology Program (RCJC20221008092720004 and 20231120201445001 to J.D.). J.D. is an investigator of the SUSTech Institute for Biological Electron Microscopy.

---

**Figure S1.**
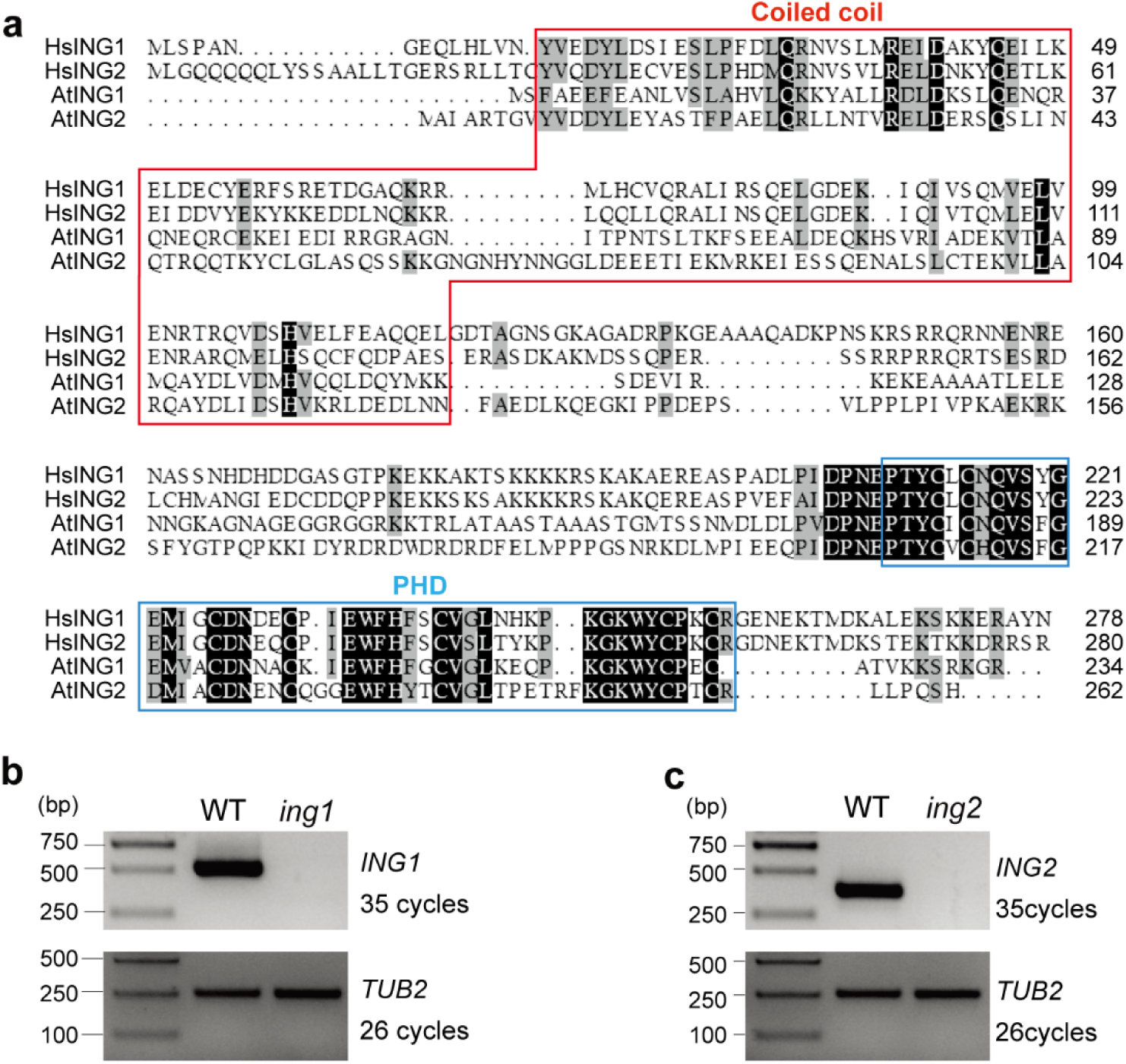
Amino acid sequence alignment of INGs and characterization of *ing1* and *ing2* mutants. **a**, Amino acid sequence alignment of *Homo sapiens* ING1 (accession: KJ891475.1), HsING2 (accession: KJ897059.1), AtING1 and AtING2. Identical residues are shaded in black, while similar residues are shaded in gray. Coiled-coil domains and PHD-finger domains are highlighted with red and blue boxes, respectively. **b**, **c**, RT-PCR analysis of *ING1* and *ING2* expression in WT, *ing1* (**b**), and *ing2* (**c**). The constitutively-expressed *TUB2* serves as an internal control.

**Figure S2.**
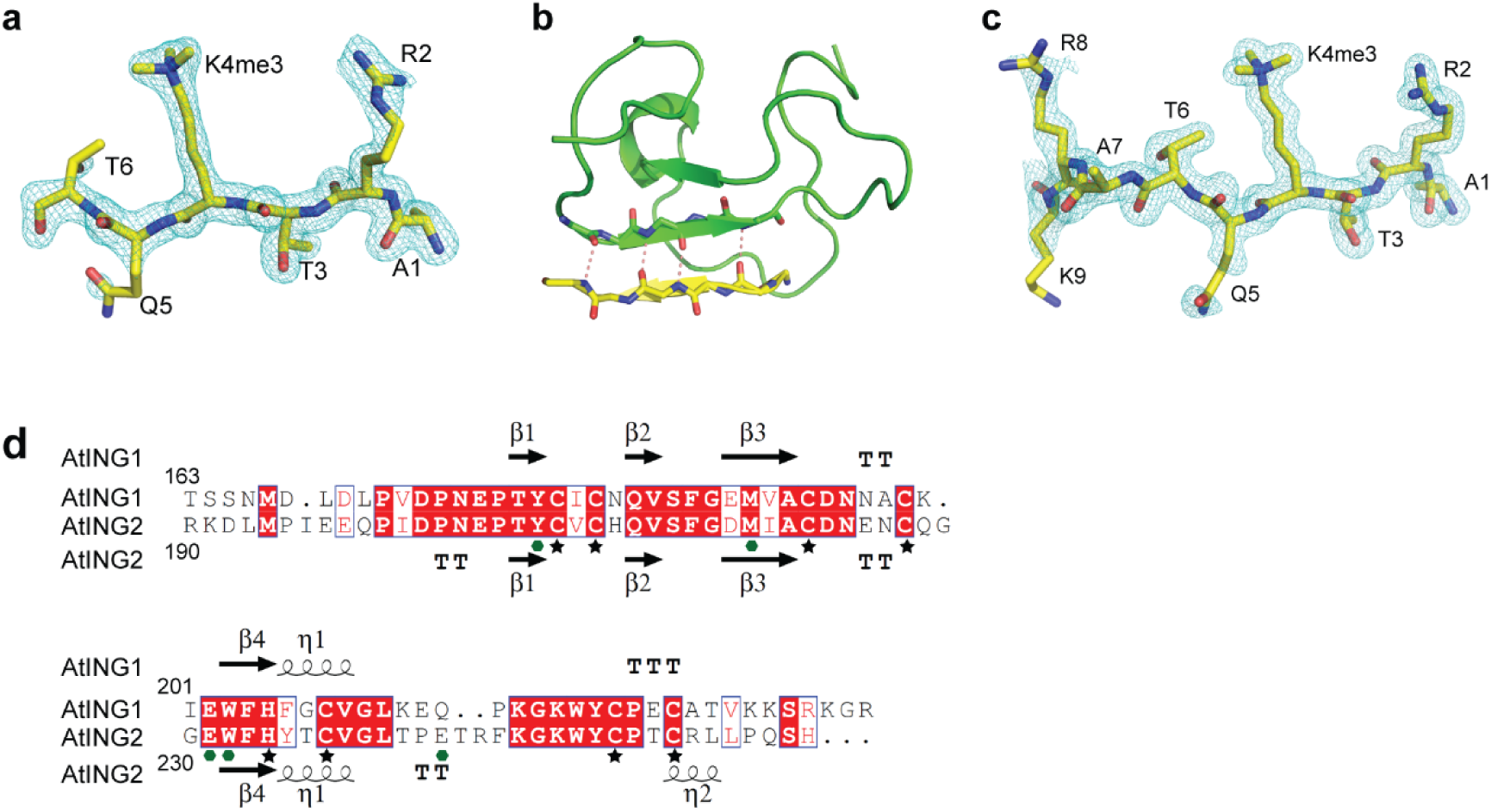
Structural analysis of the PHD-finger domains of AtING1 and AtING2 in H3K4me3 recognition. **a**, The SIGMAA weighted 2Fo-Fc electron density map of the bounded H3K4me3 peptide from the AtING1 PHD-H3K4me3 complex at 1.7 Å resolution and 1.5 sigma level. **b**, The hydrogen bonding interaction between the main chain of the peptide Arg2 to Thr6 and the main chain the PHD finger Gly189 to Ala193 are of typical antiparallel β-sheet like manner. The main chains of the two segments are shown in stick model and the hydrogen bonds are highlighted in dashed red lines. **c**, A SIGMAA weighted 2Fo-Fc electron density map of the H3K4me3 peptide from the AtING2 PHD-H3K4me3 complex at 1.6 Å resolution and 1.0 sigma level. **d**, Structure-based sequence alignment of AtING1 and AtING2 PHD fingers. The secondary structures of AtING1 and AtING2 PHD fingers are shown on the top and bottom of the alignment, respectively. The important conserved Cys and His residues to coordinate Zn²⁺ ions are marked by black stars. The conserved residues for H3K4me3 peptide binding are marked by green hexagons.

**Figure S3.**
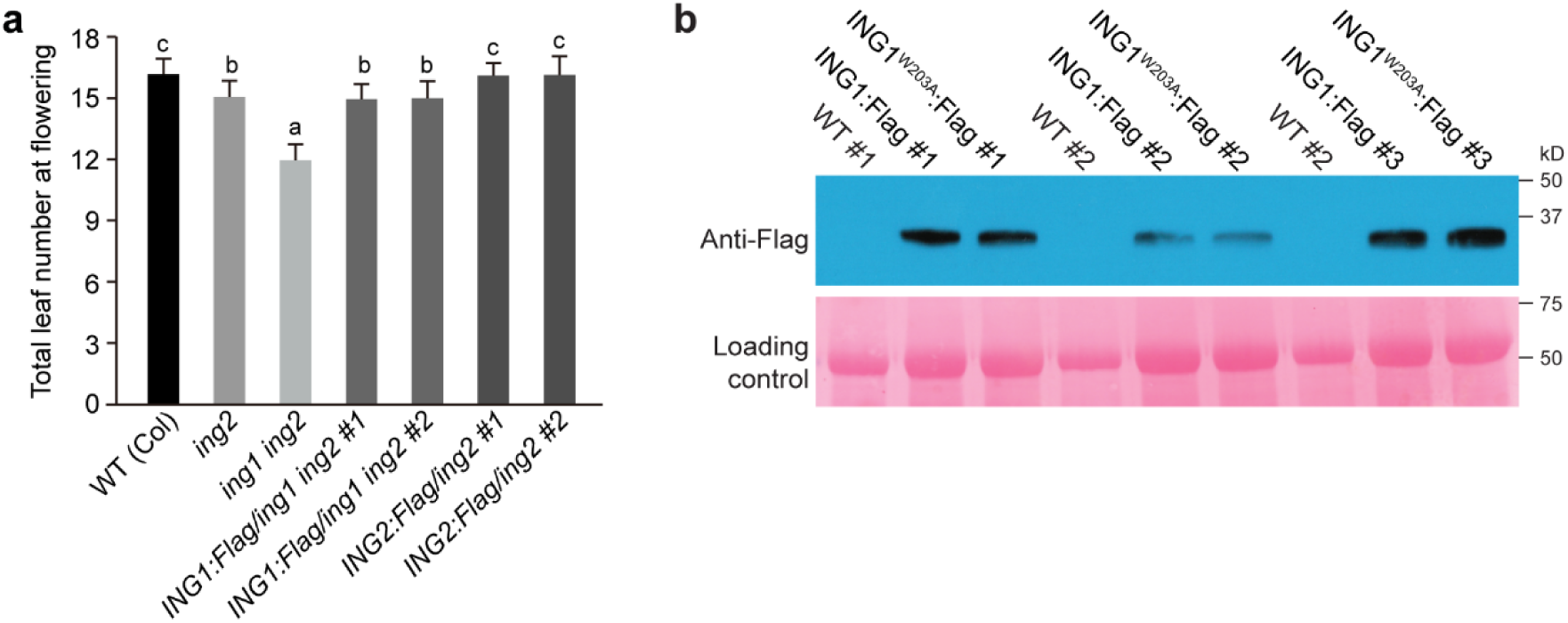
Functional analysis of *ING1/2* and protein expression of Flag-tagged ING1 variants. **a**, Analysis of *ING1pro-ING1:FLAG* and *ING2pro-ING2:FLAG* functionality. Total number of leaves at flowering was scored for each indicated line (around 20 plants per line). Error bars indicate s.d. Letters indicate statistically significant differences (one-way ANOVA, *p*<0.01). **b**, Western blot analysis to assess the abundances of ING1:FLAG and ING1^W203A^:FLAG in the indicated seedlings grown in LDs. Ponceau S-stained blot served as a loading control.

**Figure S4.**
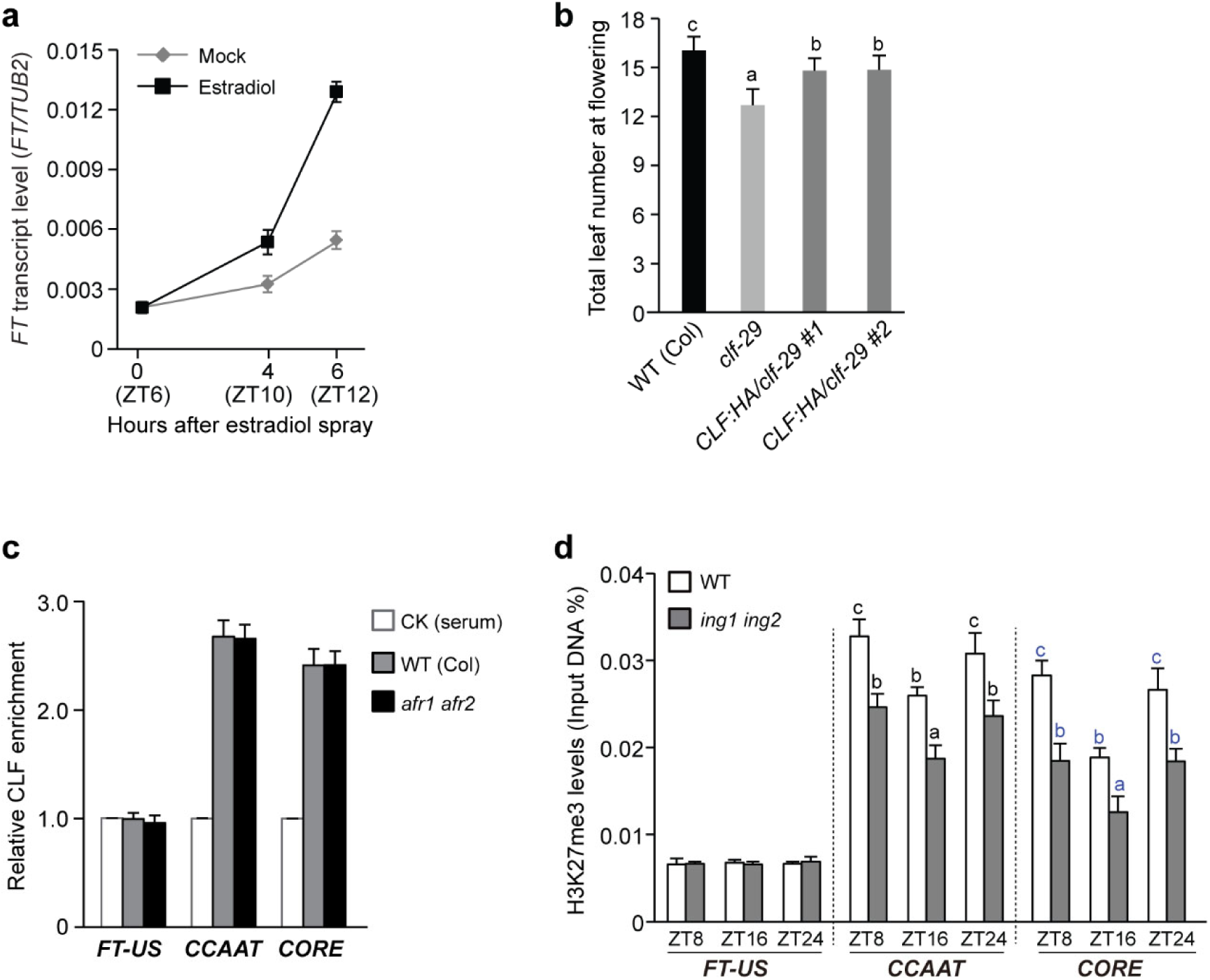
Characterization of *FT* induction as well as *FT* regulation by CLF-PRC2. **a**, *FT* induction in the estradiol-treated seedlings of doubly hemizygous *LexA_pro_-CO ING1:FLAG* grown in LDs. β-estradiol or mock was applied to the seedlings at ZT6; *FT* levels were measured at the indicated time points by RT-qPCR and directly normalized to *TUB2*. **b**, Analysis of *CLF_pro_-CLF:HA* functionality. Total number of leaves at flowering was scored for indicated lines (about 20 plants per line). Error bars indicate s.d. **c**, ChIP analysis of CLF enrichment on *FT* promoter chromatin in WT and *afr1 afr2* seedlings at ZT16 in LDs. ChIP was conducted with rabbit polyclonal anti-CLF. Relative CLF fold enrichments in each examined region over a background control (WT IP with rabbit serum), are shown. **d**, ChIP analysis of H3K27me3 on *FT* promoter chromatin in WT and *ing1 ing2* seedlings at indicated time points over a LD cycle. Levels of the genomic fragments immunoprecipitated using anti-H3K27me3 were quantified by qPCR and normalized directly to ‘input DNA’. **a-d**, Values in (**a**, **c-d**) are means ± s.d. of three biological replicates. Letters in (**b**) indicate statistically significant differences (one-way ANOVA, *p*<0.01), and letters in the same color in (**d**) indicate statistically significant differences among a group of means (one-way ANOVA, *p*<0.05).

**Table S1.**
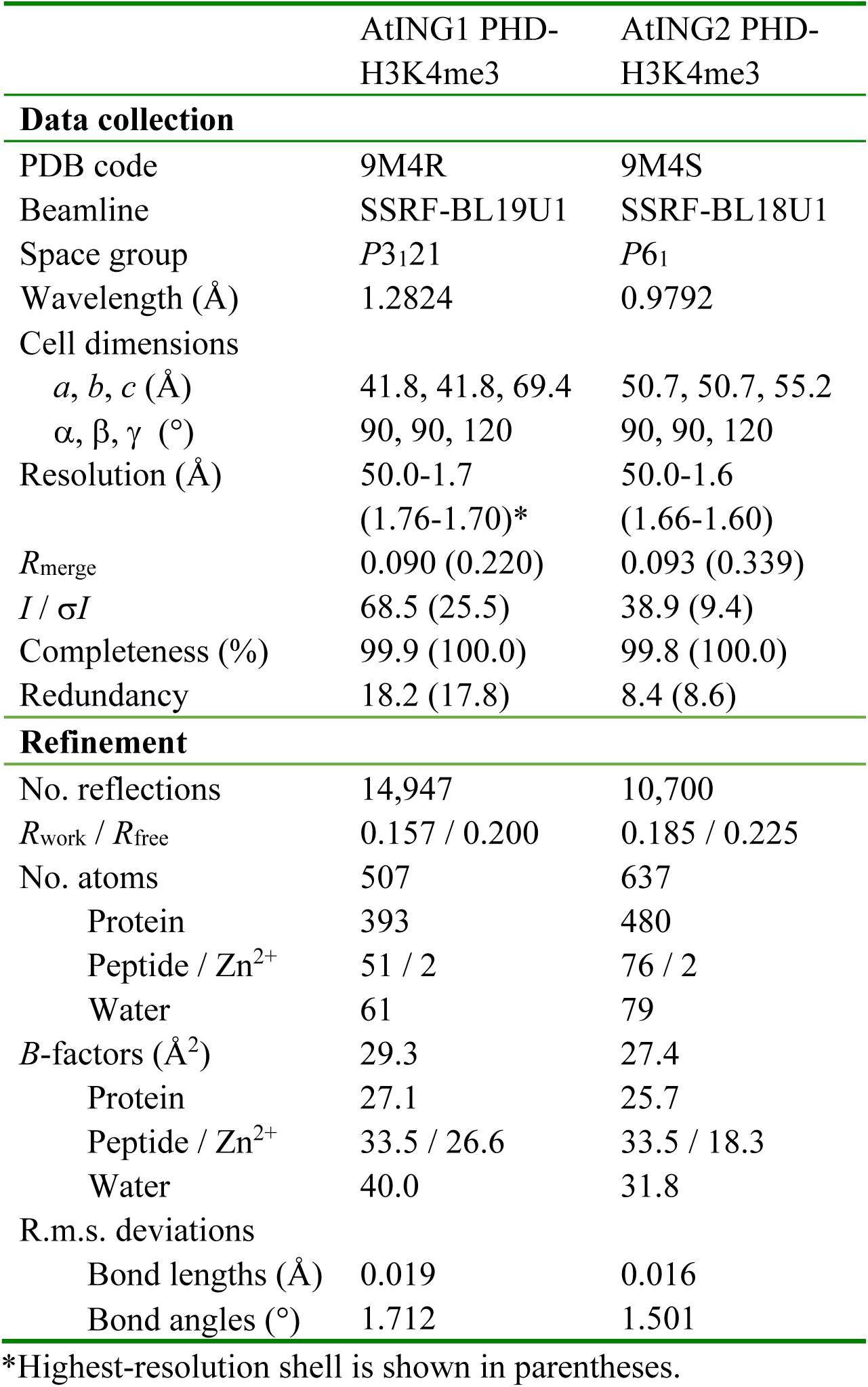
Data collection and refinement statistics for crystal structure.

|  | AtING1 PHD-H3K4me3 | AtING2 PHD-H3K4me3 |
| --- | --- | --- |
| **Data collection** |  |  |
| PDB code | 9M4R | 9M4S |
| Beamline | SSRF-BL19U1 | SSRF-BL18U1 |
| Space group | *P*3<sub>1</sub>21 | *P*6<sub>1</sub> |
| Wavelength (Å) | 1.2824 | 0.9792 |
| Cell dimensions |  |  |
| *a*, *b*, *c* (Å) | 41.8, 41.8, 69.4 | 50.7, 50.7, 55.2 |
| $\alpha$, $\beta$, $\gamma$ (°) | 90, 90, 120 | 90, 90, 120 |
| Resolution (Å) | 50.0-1.7 | 50.0-1.6 |
|  | (1.76-1.70)* | (1.66-1.60) |
| *R*<sub>merge</sub> | 0.090 (0.220) | 0.093 (0.339) |
| *I* / $\sigma I$ | 68.5 (25.5) | 38.9 (9.4) |
| Completeness (%) | 99.9 (100.0) | 99.8 (100.0) |
| Redundancy | 18.2 (17.8) | 8.4 (8.6) |
| **Refinement** |  |  |
| No. reflections | 14,947 | 10,700 |
| *R*<sub>work</sub> / *R*<sub>free</sub> | 0.157 / 0.200 | 0.185 / 0.225 |
| No. atoms | 507 | 637 |
| Protein | 393 | 480 |
| Peptide / Zn<sup>2+</sup> | 51 / 2 | 76 / 2 |
| Water | 61 | 79 |
| *B*-factors (Å<sup>2</sup>) | 29.3 | 27.4 |
| Protein | 27.1 | 25.7 |
| Peptide / Zn<sup>2+</sup> | 33.5 / 26.6 | 33.5 / 18.3 |
| Water | 40.0 | 31.8 |
| R.m.s. deviations |  |  |
| Bond lengths (Å) | 0.019 | 0.016 |
| Bond angles (°) | 1.712 | 1.501 |

\*Highest-resolution shell is shown in parentheses.

**Table S2.**
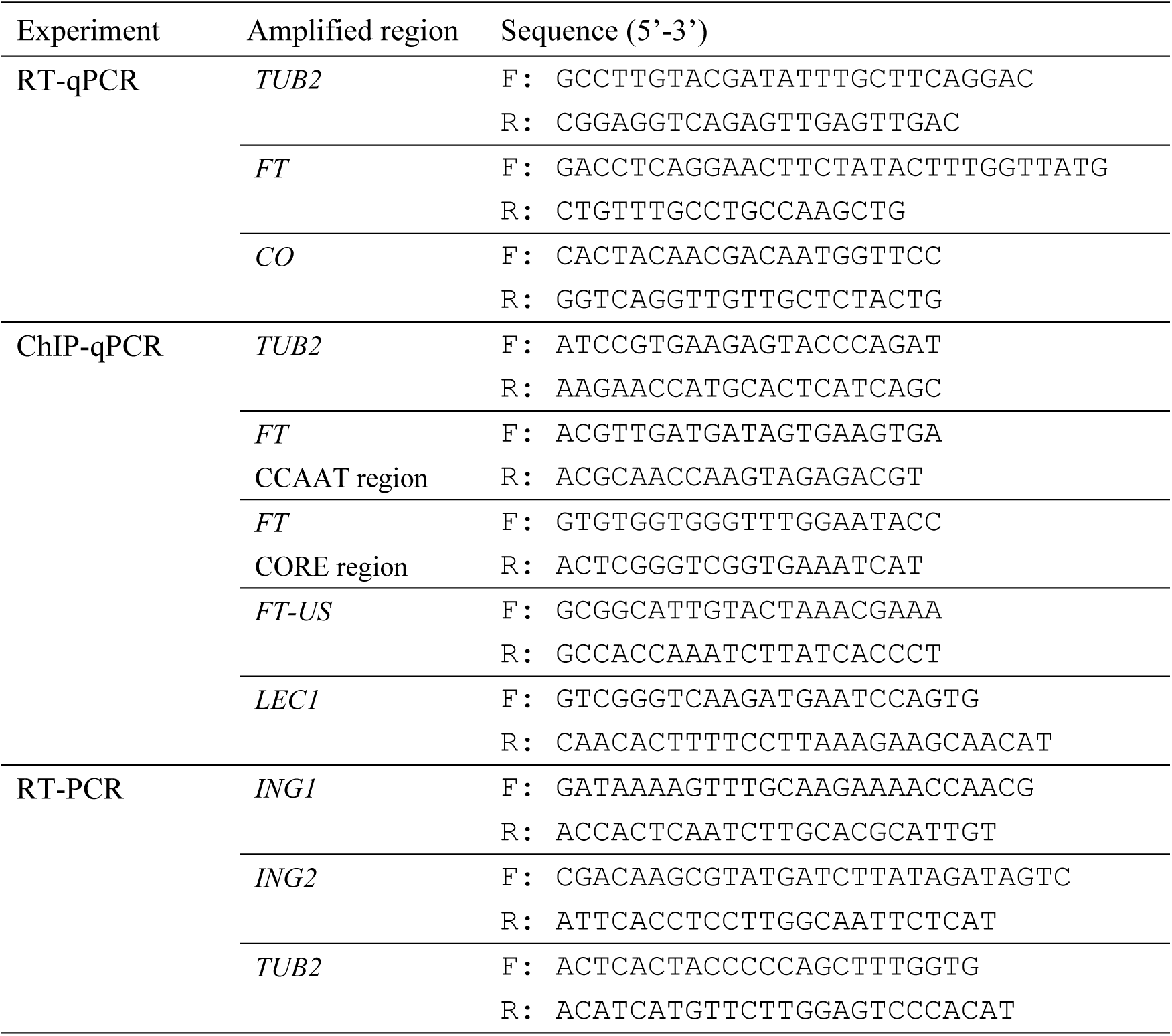
Primers used in this study.

